# NuMA Targets Dynein to Microtubule Minus-Ends at Mitosis

**DOI:** 10.1101/148692

**Authors:** Christina L. Hueschen, Samuel J. Kenny, Ke Xu, Sophie Dumont

**Author notes:** Correspondence: Sophie Dumont.

## Abstract

To build the spindle at mitosis, motors exert spatially regulated forces on microtubules. We know that dynein pulls on mammalian spindle microtubule minus-ends, and this localized activity at ends is predicted to allow dynein to cluster microtubules into poles. How dynein becomes enriched at minus-ends is not known. Here, we use quantitative imaging and laser ablation to show that NuMA targets dynactin to minus-ends, localizing dynein activity there. NuMA is recruited to new minus-ends independently of dynein and more quickly than dynactin, and both NuMA and dynactin display specific, steady-state binding at minus-ends. NuMA localization to minus-ends requires a C-terminal region outside NuMA’s canonical microtubule binding domain, and it is independent of direct minus-end binders γ-TuRC, CAMSAP1, and KANSL1/3. Both NuMA’s minus-end-binding and dynein-dynactin-binding modules are required to rescue focused, bipolar spindle organization. Thus, NuMA may serve as a mitosis-specific minus-end cargo adaptor, targeting dynein activity to minus-ends to cluster spindle microtubules into poles.

## INTRODUCTION

Each time a cell divides, the microtubule cytoskeleton remodels itself into a bipolar assembly of microtubules called the spindle. The spindle’s architecture is essential to its function – accurate chromosome segregation. In mammalian spindles, microtubule plus-ends mechanically couple to chromosomes, while microtubule minus-ends are focused into two poles that dictate where chromosomes are transported at anaphase. To build the spindle’s architecture, motors must exert spatially regulated forces on microtubules. Microtubule ends, which are structurally distinct, offer platforms for such localized regulation. Indeed, motor recruitment and activity at plus-ends is well-documented (Wu, Xiang, & Hammer, 2006), but motor regulation at minus-ends is less well understood.

Dynein is a minus-end-directed motor which slides parallel spindle microtubules to focus their minus-ends into spindle poles (Heald et al., 1996; Verde et al., 1991), working in complex with its adaptor dynactin and the microtubule-binding protein NuMA (Gaglio et al., 1996; Merdes et al., 1996). The clustering of parallel microtubules into poles presents a geometric problem when forces are indiscriminately applied all along microtubules: inversely-oriented motors between parallel microtubules will oppose each other, resulting in gridlock, unless symmetry is broken by dynein enrichment at microtubule minus-ends (Hyman & Karsenti, 1996; McIntosh, Hepler, & van Wie, 1969; Surrey et al., 2001). In computational models, localizing a minus-end-directed motor at microtubule ends permits microtubule clustering into asters or poles (Foster et al., 2015; Goshima, Nédélec, & Vale, 2005; Nedelec & Surrey, 2001; Surrey et al., 2001) and the emergence of a robust steady-state spindle length (Burbank, Mitchison, & Fisher, 2007). More recently, experimental work has shown that dynein-dynactin and NuMA do indeed selectively localize to spindle minus-ends, with dynein pulling on them after kinetochore-fiber (k-fiber) ablation in mammalian spindles (Elting et al., 2014; Sikirzhytski et al., 2014). This is consistent with suggestions that dynein and NuMA capture and pull on distal k-fiber minus-ends in monopolar spindles (Khodjakov et al., 2003). Altogether, these findings demonstrate the importance (*in silico)* and existence (*in vivo*) of localized dynein activity at spindle microtubule minus-ends.

How dynein becomes localized at minus-ends remains an open question. Dynein may be enriched near minus-ends because it walks down microtubules and piles up when it runs out of track; indeed, pile-up of dynein has been observed at minus-ends *in vitro* (McKenney et al., 2014; Soundararajan & Bullock, 2014) and can drive minus-end clustering (Tan et al., 2017). Alternatively, the exposed a-tubulin interface at microtubule minus-ends is structurally distinct and could bind an adaptor protein that specifically recruits dynein, analogous to recruitment at canonical dynein cargoes like organelles (Kardon & Vale, 2009). NuMA can target dynein-dynactin to the cell cortex (Lechler & Fuchs, 2005; Nguyen-Ngoc, Afshar, & Gonczy, 2007) and thus could be one such adaptor. However, *in vitro* NuMA has shown no direct affinity for minus-ends specifically, binding all along the microtubule lattice (Du et al., 2002; Forth et al., 2014; Haren & Merdes, 2002) or at both ends (Seldin, Muroyama, & Lechler, 2016), unlike three proteins known to interact directly with minus-ends at mitosis: γ-TuRC (Zheng et al., 1995), CAMSAP1 (Akhmanova & Hoogenraad, 2015; Hendershott & Vale, 2014; Jiang et al., 2014) and KANSL1/3 (Meunier et al., 2015). In cells, NuMA is thought to require dynein activity to carry it to minus-ends and spindle poles (Merdes et al., 2000), where it anchors spindle microtubules (Dionne, Howard, & Compton, 1999; Gaglio, Saredi, & Compton, 1995; Heald et al., 1997; Silk, Holland, & Cleveland, 2009). Thus, it remains unclear whether dynein-dynactin and NuMA have specific binding sites at minus-ends, and if so, whether they are recruited by known minus-end binders. Finally, knowing how dynein is targeted to minus-ends would make it possible to test the *in vivo* role of minus-end-localized – compared to indiscriminately-localized –forces in spindle organization.

Here, we use laser ablation to create new, isolated minus-ends in the mammalian spindle *in vivo*, and quantitative imaging to map protein recruitment to these ends and its mechanistic basis. We demonstrate that NuMA binds at minus-ends independently of dynein, and that NuMA targets dynactin – and thereby dynein activity – to spindle minus-ends. This challenges the prevailing model that dynein delivers NuMA to spindle minus-ends and poles (Merdes et al., 2000; Radulescu & Cleveland, 2010). NuMA localization to minus-ends is independent of known direct minus-end binders γ-TuRC, CAMSAP1, and KANSL1/3, and it requires both NuMA’s canonical microtubule-binding domain and an additional region of its C-terminus. Thus, NuMA –which is sequestered in the nucleus at interphase (Lydersen & Pettijohn, 1980) – may serve as a mitosis-specific minus-end cargo adaptor, recruiting dynein activity to spindle minus-ends. Both NuMA’s minus-end-binding domain and dynactin-binding domain are required for correct spindle architecture, supporting long-standing *in silico* predictions that localizing dynein to minus-ends enables effective clustering of parallel microtubules into poles. These findings identify a mechanism for mitosis-specific recruitment of dynein to microtubule minus-ends and, more broadly, illustrate how spatial regulation of local forces may give rise to larger-scale cytoskeletal architectures.

## RESULTS

### Dynactin and NuMA display mitosis-specific, steady-state binding at microtubule minus-ends

To visualize the spatial targeting of the dynein-dynactin complex to microtubule minus-ends – which are normally buried in dense mammalian spindles – we used nocodazole washout and laser ablation to create resolvable minus-ends in mitotic cells. First, to determine whether dynein-dynactin and NuMA localize to individual microtubule minus-ends, we treated mammalian PtK2 and RPE1 cells with the microtubule-depolymerizing drug nocodazole and fixed cells 6-8 min after drug washout to capture acentrosomal microtubules with clearly visible plus- and minus-ends (Figure 1A). p150*^Glued^* (p150, a dynactin subunit) and NuMA strongly co-localized at the minus-ends of these individual microtubules, with a clear binding preference for minus-ends over the microtubule lattice or the plus-end, which is marked by EB1 (Figure 1B). Interestingly, in prophase cells before nuclear envelope breakdown, p150 localized predominantly to plus-ends rather than minus-ends (Figure 1B; Figure 1-figure supplement 1), consistent with dynactin’s interphase localization (Vaughan et al., 1999). Thus, nuclear envelope breakdown (NEB) confers dynactin’s preference for minus-ends, suggesting regulated, mitosis-specific spatial targeting.

**Figure 1.**
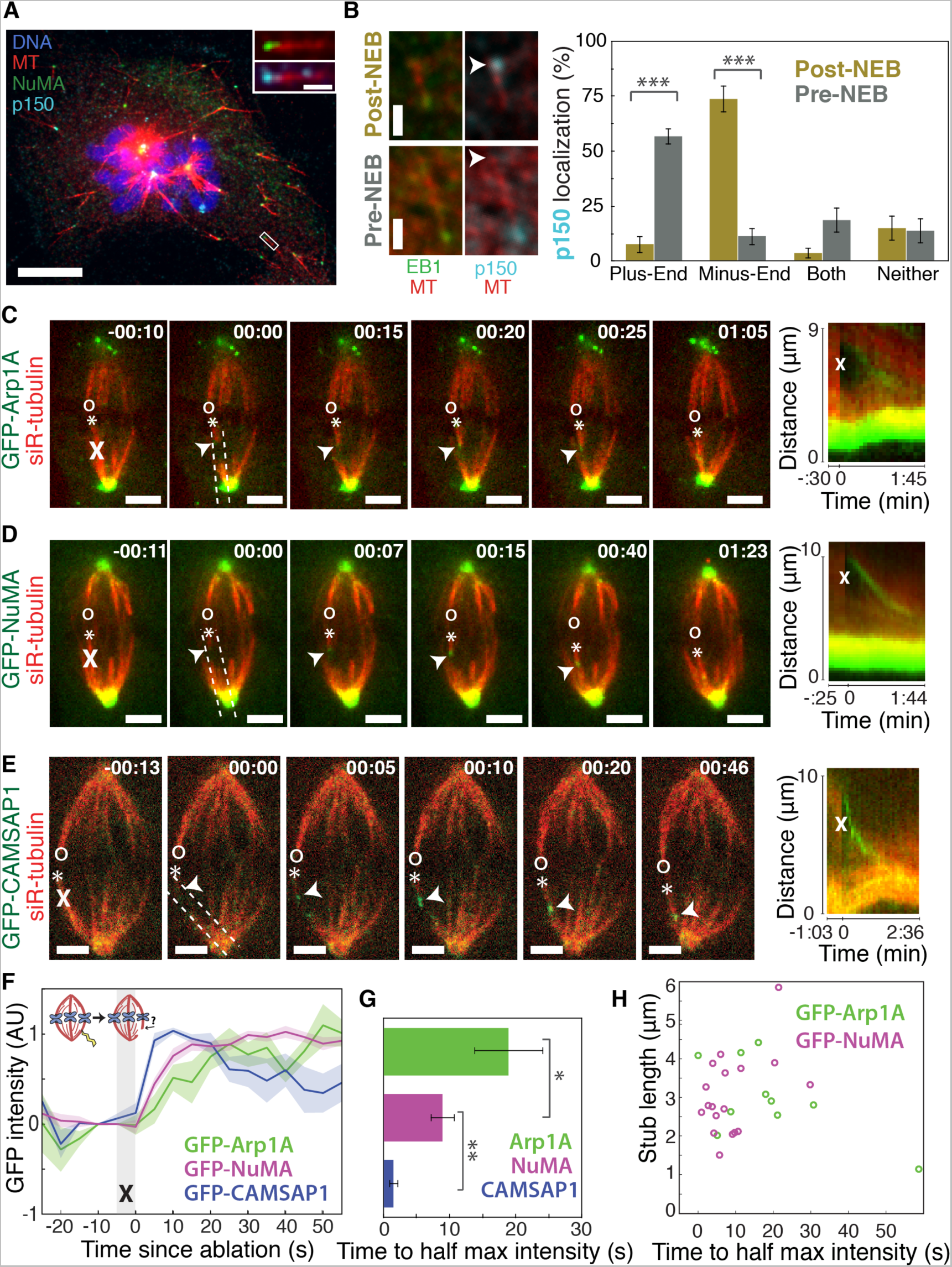
Dynactin and NuMA display specific, steady-state binding at mitotic minus-ends. See also Figure 1-figure supplement 1 and Videos 1-3. (A) Representative immunofluorescence image showing co-localization of NuMA (green) and p150 (dynactin subunit; cyan) at microtubule minus-ends in mitotic PtK2 cells (post-NEB) fixed after washout of 5 μM nocodazole. Scale bar, 10 μm. Inset: zoom of white box, with 1 μm scale bar. (B) Representative immunofluorescence images of mitotic RPE1 cells, processed as in (A). After nuclear envelope breakdown (post-NEB), EB1 (green) and p150 (cyan) localize to opposite microtubule ends. In prophase cells (pre-NEB), p150 instead co-localizes with EB1 at plus-ends. Scale bar, 1 μm. Graph displays mean percentage ± SEM of p150 at each location within one cell for *n* = 57 microtubules, 16 cells (post-NEB); *n* = 72 microtubules, 8 cells (pre-NEB). \*\*\**P* < 0.0001. (C-E) Representative time-lapse live images and kymographs of PtK2 cells expressing (C) GFP-Arp1A, (D) GFP-NuMA, or (E) GFP-CAMSAP1. Arp1A, NuMA, and CAMSAP1 are all recruited to k-fiber minus-ends (arrowheads) created by ablation (‘X’s) and move with them as dynein pulls them poleward (Elting et al., 2014). Time is in min:sec, with the frame captured immediately following ablation set to 00:00. ‘*’ marks plus-end of ablated k-fiber, and ‘o’ marks its sister. Scale bar, 5 μm. Kymographs (right) are along poleward paths (dashed lines). (F) Plot of mean normalized GFP intensity and SEM (shading) over time at ablation-created minus-ends for GFP-Arp1A, GFP-NuMA, and GFP-CAMSAP1. Time = 0 s at the first frame following ablation. *n* =10 ablations, 7 cells (Arp1A); *n* =18 ablations, 7 cells (NuMA); *n* =13 ablations, 7 cells (CAMSAP1). (G) Time from ablation to half maximum GFP intensity, calculated for each individual ablation (see Methods) and then averaged for data in (F). Error bars show SEM. \**P* < 0.05; \*\**P* < 0.01. (H) Time to half-maximum GFP-Arp1A or GFP-NuMA intensity at ablation-created minus-ends as a function of length of ablation-created k-fiber stubs. Correlation coefficient = −0.62, *P* = 0.06, *n* = 10 (Arp1A); correlation coefficient = 0.44, *P* = 0.07, *n* = 18 (NuMA).

Second, we sought to test whether dynactin and NuMA have finite binding sites at minus-ends by measuring the kinetics of their recruitment. To do so, we used laser ablation of k-fibers in PtK2 cells to create new minus-ends within the spindle body (Figure 1C-E). By spatially and temporally synchronizing the creation of a bundle of minus-ends, laser ablation allowed for dynamic measurements of the recruitment to minus-ends of GFP-tagged dynactin (Arp1A) and NuMA and comparison to a direct minus-end binding protein, CAMSAP1. Dynactin, NuMA, and CAMSAP1 robustly recognized new microtubule minus-ends within the spindle (Figure 1C-E). The kinetics of dynactin and NuMA recruitment to minus-ends were distinct from CAMSAP1, which reached a max intensity after approximately 5-10 s, but then decreased in intensity as if its binding sites at the minus-end were gradually obstructed (Figure 1E-G). Unlike CAMSAP1, dynactin and NuMA intensities reached a stable plateau, suggesting that binding saturates, reaching a steady-state. This finite binding is consistent with a truly end-specific localization (rather than localization along the unbounded lattice *near* the minus-end). In addition, the rate of NuMA or dynactin accumulation at new minus-ends did not correlate with the length of k-fiber stubs created by ablation (Figure 1H), which could indicate that recruitment rate is set by the number of individual microtubule minus-ends (which is similar across k-fibers (McEwen et al., 1997)) rather than k-fiber length.

Interestingly, NuMA intensity at new minus-ends increased at a faster rate and saturated sooner than dynactin (Figure 1F,G). This observation hints at a model in which NuMA targets dynein-dynactin to minus-ends, and it is less easily consistent with the idea that dynein delivers NuMA to minus-ends. If true, this hypothesis would explain why dynactin’s minus-end preference arises after NEB, when NuMA is released from the nucleus into the mitotic cytoplasm.

### NuMA localizes to minus-ends independently of dynein

The finding that NuMA localizes to minus-ends more quickly than dynactin could suggest that NuMA directly or indirectly recognizes the exposed α-tubulin structure of minus-ends and subsequently recruits dynein-dynactin (‘Structural Recognition,’ Figure 2A). Alternatively, dynein-dynactin could walk toward minus-ends, carrying NuMA, and pile up or dwell there (‘Walk and Pile-up,’ Figure 2B). To test the ‘Walk and Pile-up’ model, we inhibited dynein-dynactin in PtK2 spindles by over-expressing p50 (dynamitin), resulting in fully unfocused k-fiber arrays (Figure 2C). The ‘Walk and Pile-up’ model predicts that in the absence of dynein-dynactin transport, NuMA – which requires dynactin for its interaction with dynein (Merdes et al., 2000) – should not reach minus-ends. Instead, GFP-NuMA was robustly recruited to minus-ends created by k-fiber ablation (Figure 2C), indicating that NuMA can localize to minus-ends independently of dynein. Similarly, NuMA was recruited to ablation-created minus-ends when dynein was inhibited by p150-CC1 over-expression (Figure 2-figure supplement 1A,B). Thus, NuMA localizes to minus-ends without dynein carrying it there – consistent with early observations in extract asters and spindles (Gaglio et al., 1996; Heald et al., 1997) but in contrast to the prevailing view that dynein delivers NuMA to minus-ends (Merdes et al., 2000; Radulescu & Cleveland, 2010). Slower NuMA accumulation kinetics after p50 overexpression suggest that dynein-dynactin-NuMA complex formation may aid rapid NuMA recruitment, but dynein ‘Walk and Pile-up’ alone cannot explain NuMA’s minus-end affinity (Figure 2D,E).

**Figure 2.**
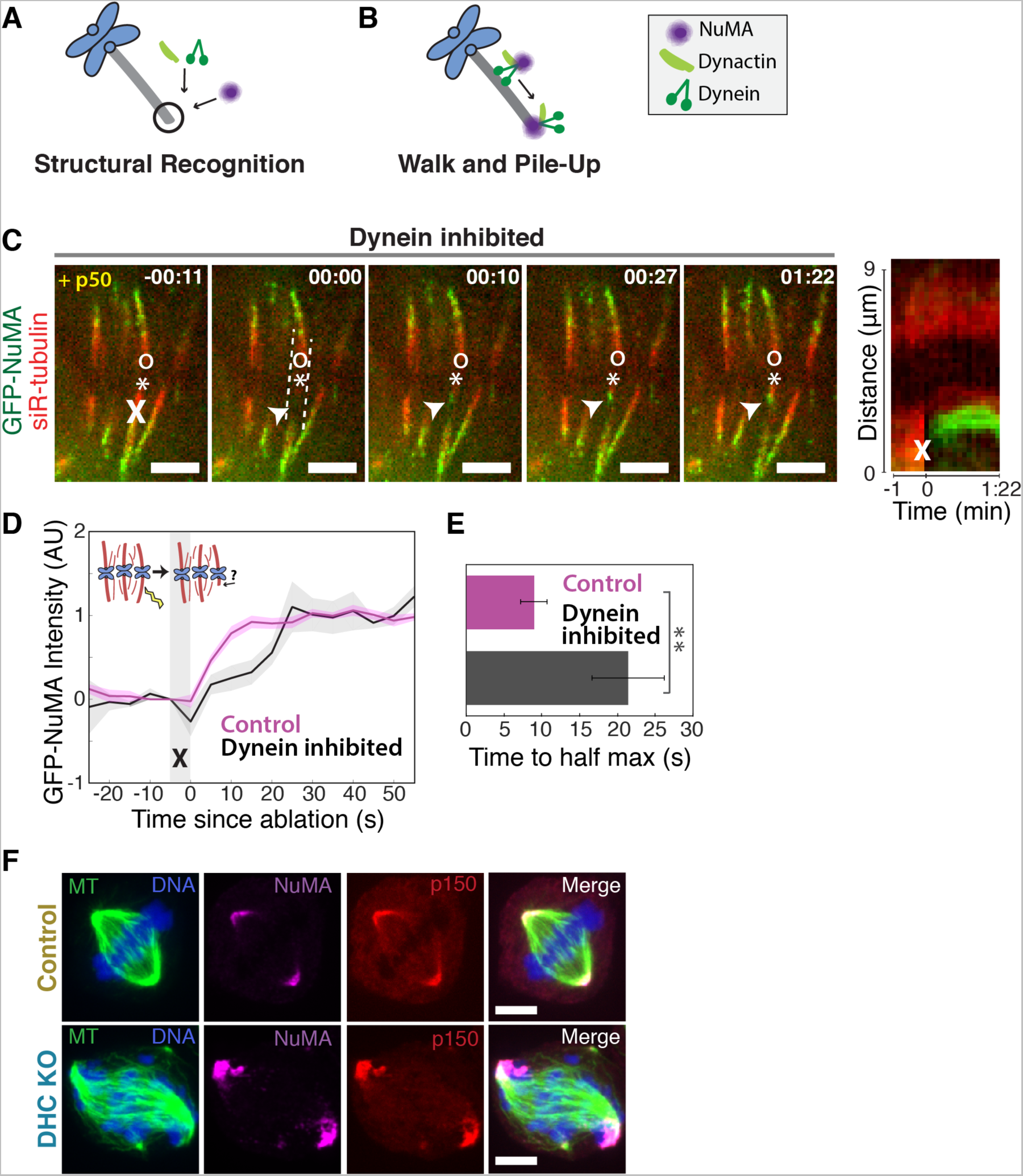
NuMA can localize to minus-ends independently of dynein. See also Figure 2-figure supplement 1 and Video 4. (A-B) Models for targeting dynein to minus-ends. Model A, Structural Recognition: Cytoplasmic molecules may recognize the exposed a-tubulin interface at minus-ends and recruit dynein. Model B, Walk and Pile-up: Minus-end-directed dynein may walk to minus-ends and pile up. (C) Representative time-lapse live images and kymograph of PtK2 cell expressing GFP-NuMA, in which dynein-dynactin is inhibited by overexpression of p50 (dynamitin) (Shrum et al., 2009). K-fibers are unfocused and splayed, but NuMA is still robustly recruited to k-fiber minus-ends (arrowheads) created by ablation (‘X’). Time is in min:sec, with frame captured immediately following ablation set to 00:00. ‘*’ marks plus-end of ablated k-fiber, and ‘o’ marks its sister. Scale bars, 5 μm. Kymograph (right) taken along dashed line path. (D) Plot of mean normalized GFP-NuMA intensity and SEM (shading) over time at ablation-created minus-ends. Time = 0 s at the first frame following ablation. *n* = 18 ablations, 7 cells for control; *n* = 14 ablations, 8 cells for dynein-inhibited (p50 overexpression). \*\**P* = 0.01. (E) Time from ablation to half maximum GFP-NuMA intensity, calculated for each individual ablation (see Methods) and then averaged for data in (D). Error bars show SEM. (F) Representative immunofluorescence images of inducible-Cas9 dynein heavy chain (DHC)-knockout HeLa cells (McKinley & Cheeseman, 2017) showing robust localization of NuMA and p150 (dynactin) at minus-ends when (DHC) is deleted. Scale bar, 5 µm.

To confirm that NuMA localizes to minus-ends even after genetic dynein deletion, we used an inducible CRISPR/Cas9 HeLa cell line to knock out dynein’s heavy chain (DHC) (McKinley & Cheeseman, 2017). After dynein knockout (Figure 2-figure supplement 1C), both NuMA and dynactin localized robustly to k-fiber minus-ends (Figure 2F). Together, these data indicate that NuMA can bind at minus-ends in the absence of dynein and are consistent with NuMA-mediated recognition of minus-end structure (‘Structural Recognition,’ Figure 2A). In addition, they suggest that NuMA may interact with and target dynactin to minus-ends even in the absence of dynein, when dynein-dynactin complexes cannot form and dynein motility cannot deliver dynactin close to minus-ends.

### NuMA is required for dynein activity and dynactin localization at minus-ends

If NuMA does target dynein-dynactin to minus-ends, localizing force there, we would expect NuMA to be required for dynein to transport minus-ends created by k-fiber ablation (Elting et al., 2014; Sikirzhytski et al., 2014). To test this hypothesis, we made inducible CRISPR/Cas9 RPE1 cell lines to knock out NuMA (Figure 3-figure supplement 1) (McKinley et al., 2015; T. Wang et al., 2015). Indeed, when NuMA was knocked out – causing elongated, heterogeneously disorganized spindles with detached centrosomes – ablation-created minus-ends were no longer transported toward poles by dynein (Figure 3A-C). These data are consistent with a previous finding that NuMA antibody injection prevents distal k-fiber looping events in monopolar spindles, in which dynein likely pulls on free k-fiber minus-ends (Khodjakov et al., 2003). Thus, NuMA is required for dynein activity at spindle microtubule minus-ends.

**Figure 3.**
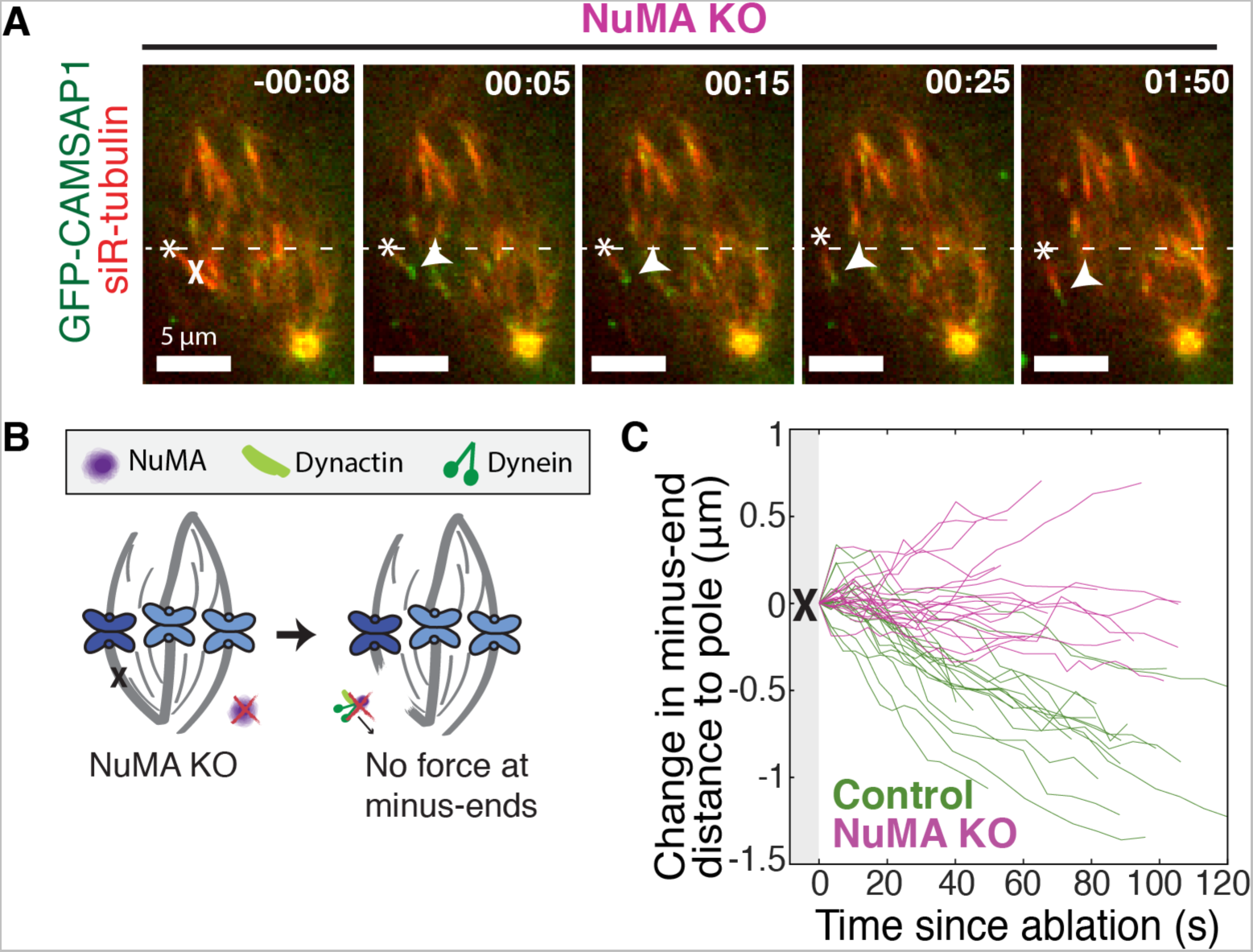
NuMA is required for force generation at minus-ends. See also Figure 3-figure supplement 1 and Video 5. (A) Representative time-lapse live images and kymograph of RPE1 cell in which NuMA has been knocked out using an inducible Cas9 system. GFP-CAMSAP1 is expressed to label minus-ends. After ablation (‘X’), the k-fiber minus-end (arrowhead) is not transported poleward by dynein and remains detached from the spindle. Time is in min:sec, with the frame captured immediately following ablation set to 00:00 s. Plus-end of ablated k-fiber is marked by ‘*’. Scale bars, 5 μm. Kymograph (right) taken along dashed line path. (B) Cartoon: In the absence of NuMA, force generation at ablation-created minus-ends is not observed. (C) Movement of ablation-created minus-ends (marked by GFP-CAMSAP1) relative to the pole. In control cells (green traces), minus-ends are transported toward the pole by dynein at consistent speeds, but this transport is lost when NuMA is knocked out (purple traces). *n* =19 ablations, 8 cells (control); *n* = 20 ablations, 6 cells (NuMA knockout).

Dynein activity at minus-ends could require NuMA because NuMA modulates dynein-dynactin’s ability to pull on microtubules, or because NuMA localizes dynein to minus-ends. Given the findings in Figures 1-2, we hypothesized that NuMA localizes dynein activity by recruiting dynactin to minus-ends. Indeed, after NuMA knockout, dynactin (GFP-Arp1A) was no longer detectable at k-fiber minus-ends created by laser ablation (Figure 4A). Immunofluorescence experiments confirmed that in the absence of NuMA, dynactin remained at kinetochores but no longer localized to minus-ends (Figure 4B). Interestingly, dynein’s localization within the spindle (labeled using an antibody against dynein intermediate chain) was less minus-end-specific than dynactin’s, both before and after NuMA deletion (Figure 4-figure supplement 1; Figure 4B).

**Figure 4.**
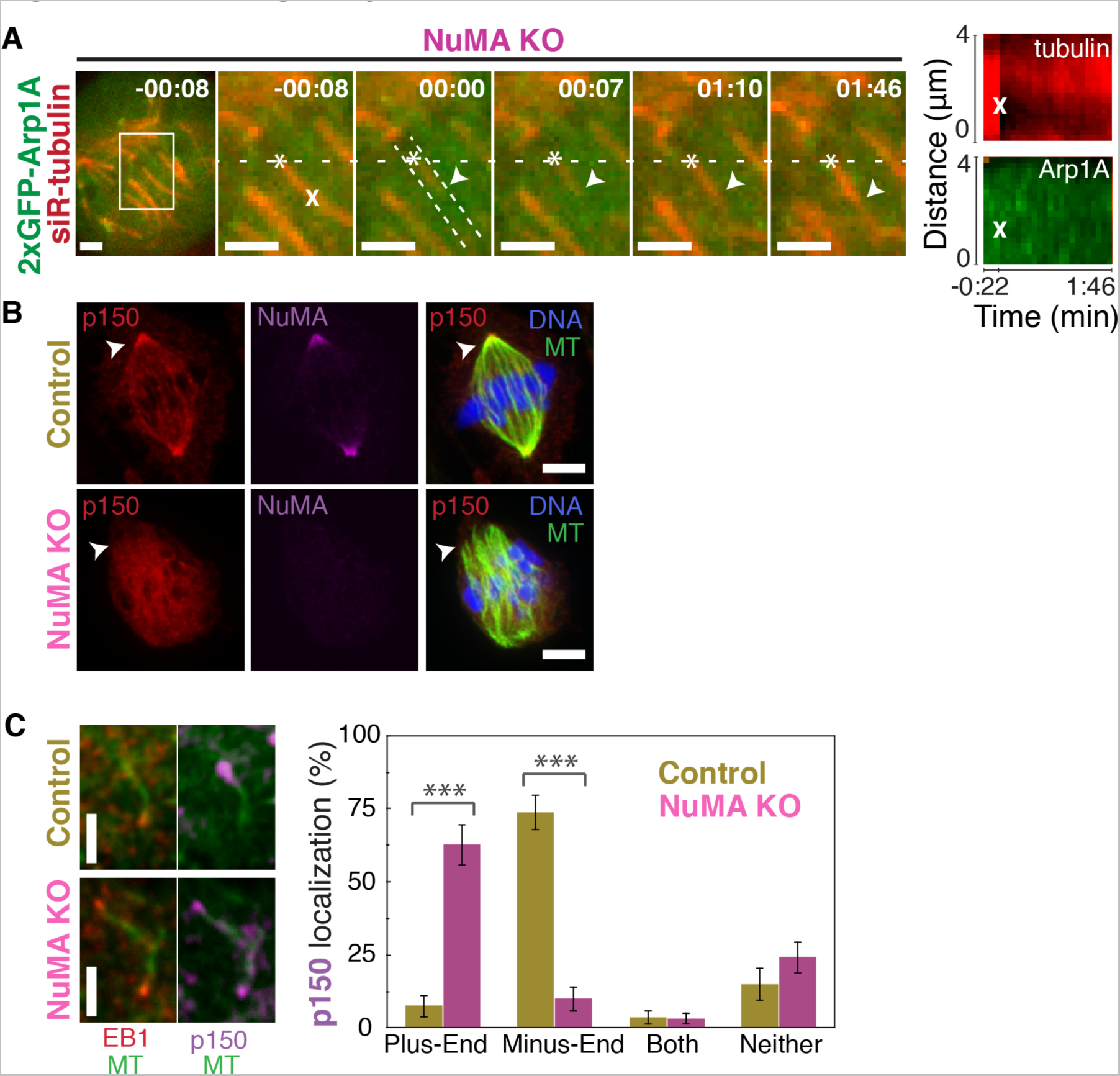
NuMA targets dynactin to mitotic minus-ends. See also Figure 4-figure supplement 1 and Video 6. (A) Representative time-lapse live images and kymograph of RPE1 cell in which NuMA has been knocked out using an inducible Cas9 system. 2xGFP-Arp1A recruitment was never observable at minus-ends (arrowheads) created by ablation (‘X’) (*n* =16 ablations, 6 cells). Time is in min:sec, with frame captured immediately following ablation set to 00:00. Plus-end of ablated k-fiber is marked by ‘*’. Scale bars, 2 μm. Kymographs (right) taken along dashed line path. (B) Representative immunofluorescence images of control and NuMA-knockout RPE1 spindles, showing a loss of p150 (dynactin) at minus-ends in the absence of NuMA. Scale bars, 5 μm. (C) Representative immunofluorescence images of mitotic RPE1 cells (post-NEB) fixed after washout of 5 μM nocodazole, as in Figure 1A,B. In control cells, p150 (dynactin; purple) localizes to microtubule minus-ends, opposite EB1 (red). When NuMA is knocked out, p150 co-localizes with EB1 at plus-ends. Scale bars, 2 μm. Graph displays mean percentage ± SEM of p150 at each location within each cell, for *n* = 72 microtubules, 8 cells (control); *n* = 72 microtubules, 11 cells (NuMA knockout). \*\*\**P* < 0.0001.

In addition, nocodazole washout experiments revealed that dynactin (p150) localization to individual microtubule minus-ends was lost upon NuMA deletion (Figure 4C). Instead, dynactin frequently co-localized with EB1 at plus-ends, similar to its interphase and pre-NEB localization pattern (Figure 4C; Figure 1B). Thus, the data indicate that NuMA is required for the transport of minus-ends by dynein because NuMA binds at minus-ends and recruits dynactin there.

### NuMA recognizes minus-ends independently of known minus-end binders

How is NuMA targeted to minus-ends, independently of dynein? *In vitro*, canonical microtubule-binding regions of NuMA have shown no preference for minus-ends relative to the lattice or plus-end of purified microtubules (Du et al., 2002; Forth et al., 2014; Haren & Merdes, 2002; Seldin et al., 2016). Given this lack of minus-end-specific binding *in vitro*, we hypothesized that NuMA is indirectly recruited to spindle minus-ends by one of three known direct minus-end binders active at mitosis: γ-TuRC (Zheng et al., 1995), CAMSAP1 (Hendershott & Vale, 2014; Jiang et al., 2014) (Figure 1; CAMSAP2 and 3 are phosphorylated at mitosis and no longer interact with microtubules (Jiang et al., 2014)), and KANSL1/3 (Meunier et al., 2015). To test this hypothesis, we treated RPE1 cells with 30 μM gatastatin to block γ-TuRC binding (Chinen et al., 2015), resulting in short spindles with fewer astral microtubules (Figure 5A; Figure 5-figure supplement 1A). We also made inducible CRISPR/Cas9 RPE1 cell lines to knock out CAMSAP1 or KANSL1 (Figure 5-figure supplement 1B,C), as KANSL1 depletion has been shown to delocalize the entire KANSL complex (Meunier et al., 2015). CAMSAP1 knockout caused a small reduction in spindle length (Figure 5-figure supplement 1D), consistent with the spindle minus-end protecting function ascribed to its *Drosophila* homolog, Patronin (Goodwin & Vale, 2010). To our surprise, however, none of these perturbations qualitatively altered NuMA localization at spindle poles (Figure 5A). To check for a subtler contribution of γ-TuRC, CAMSAP1, or KANSL1 to NuMA localization at minus-ends, we performed k-fiber ablations after 30 μM gatastatin treatment, CAMSAP1 knockout, or KANSL1 knockout and quantified GFP-NuMA recruitment kinetics at new minus-ends. NuMA recruitment to new minus-ends remained robust, and recruitment timescales were statistically indistinguishable from control (Figure 5B-C). Thus, the data indicate that the direct mitotic minus-end binders γ-TuRC, CAMSAP1, and KANSL1/3 are not responsible for NuMA’s localization to spindle microtubule minus-ends.

**Figure 5.**
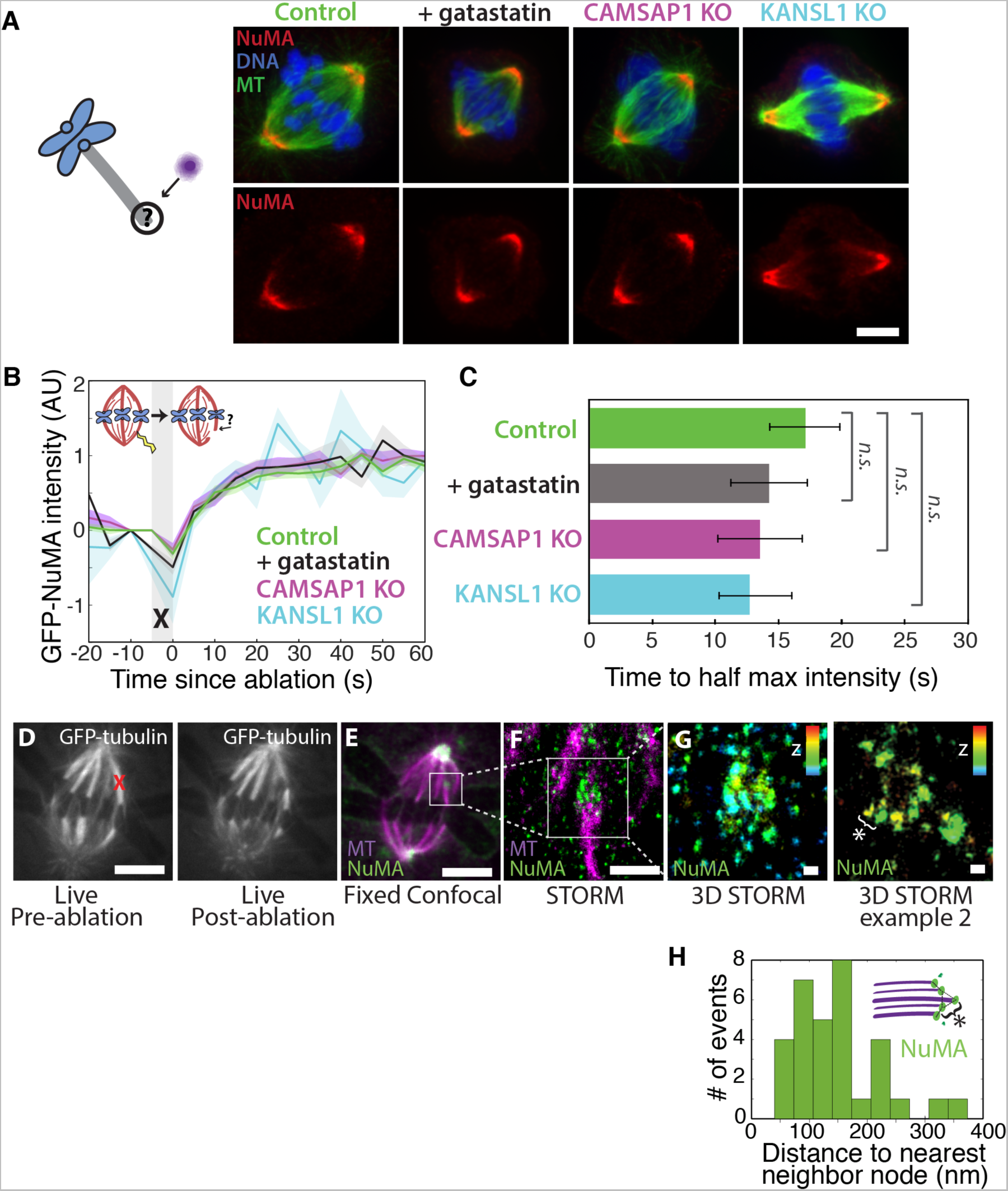
NuMA localizes to minus-ends independently of known minus-end binding proteins. See also Figure 5-figure supplement 1. (A) Schematic of hypothesis that a minus-end binding protein recruits NuMA. Instead, representative immunofluorescence images show unchanged NuMA localization in control RPE1 cells and RPE1 cells in which direct mitotic minus-end binders are inhibited (30 μM gatastatin to inhibit γ-tubulin) or knocked out (CAMSAP1, KANSL1). Scale bar, 5 μm. (B) Plot of mean normalized GFP-NuMA intensity and SEM (shading) over time at ablation-created minus-ends. Time = 0 s at the first frame following ablation. *n* = 18 ablations, 7 cells (control); *n* = 9 ablations, 8 cells (+gatastatin); *n* =10 ablations, 5 cells (CAMSAP1 knockout); *n* = 8 ablations, 4 cells (KANSL1 knockout). ‘*n.s.*’ indicates *P* > 0.05. (C) Time from ablation to half maximum GFP-NuMA intensity was calculated for each individual ablation (see Methods) and then averaged for data in (B). Error bars show SEM. (D-G) 3D STORM of NuMA at PtK2 k-fiber minus-ends created by ablation. (D) K-fibers were cut using ablation (red ‘X’), and cells were immediately fixed for immunofluorescence. Individual ablated spindles were imaged by spinning-disk confocal (E) and then by 3D STORM (F,G). In two right-most panels (G), structures are colored according to position in the Z-axis (red = + 300 nm, blue = −300 nm). Scale bars: 5 μm in (D,E); 1 μm in (F); 100 nm in (G). (E) Distances between neighboring ‘nodes’ of NuMA (e.g., ‘*’ here and in (D)) are ~50-150 nm, consistent with measured spacing between individual microtubules within PtK2 k-fibers (McDonald et al., 1992). *n* = 32 nodes, 4 ablations.

Given this lack of involvement of direct minus-end binding proteins, we sought to confirm using super-resolution microscopy that NuMA specifically localizes at individual spindle microtubule minus-ends. Three-dimensional stochastic optical reconstruction microscopy (3D STORM (Huang et al., 2008; Rust, Bates, & Zhuang, 2006)) of NuMA at k-fiber minus-ends created by ablation revealed organized clusters of NuMA puncta, rather than a lawn of molecules along the lattice near the minus-end (Figure 5D). The spacing between puncta was consistent with the distance between individual microtubules within k-fibers (Figure 5E), as measured by electron microscopy in the same cell type (PtK2) (McDonald et al., 1992), consistent with the idea that these NuMA puncta are built on individual microtubule minus-ends. A yet undiscovered minus-end binding protein may recruit NuMA; alternatively, NuMA may have direct minus-end-specific binding ability that has not been recapitulated *in vitro*.

### NuMA function requires both minus-end-recognition and dynactin-recruitment modules

To define the NuMA domain required for spindle minus-end localization, we performed rescue experiments with different NuMA truncations in the NuMA-knockout background (Figure 6A). This avoids the C-terminus-mediated oligomerization (Harborth et al., 1999) with endogenous protein that has complicated interpretations of previous work. In this NuMA-knockout background, full-length NuMA (‘FL’) localized to spindle minus-ends, as did a bonsai construct (‘N-C’) with most of the central coiled-coil removed (Figure 6B). To our surprise, ‘N-C’ rescued spindle architecture as effectively as full-length NuMA, indicating that the extraordinary length of NuMA protein is not essential to its function in spindle structure (Figure 6C). Importantly, NuMA’s C-terminus (‘C’) alone localized to minus-ends even in the absence of endogenous full-length protein (Figure 6B). Its intensity at minus-ends was less striking than that of full-length or N-C protein at poles, likely because minus-ends are not as densely concentrated in disorganized spindles, and perhaps NuMA’s N-terminus facilitates formation of higher order NuMA assemblies. However, the preference of NuMA ‘C’ for spindle ends was clear. Because dynein-dynactin interacts with NuMA’s N-terminus (Kotak, Busso, & Gönczy, 2012), the minus-end localization of NuMA’s C-terminus provides further support for dyneinindependent minus-end recognition.

**Figure 6.**
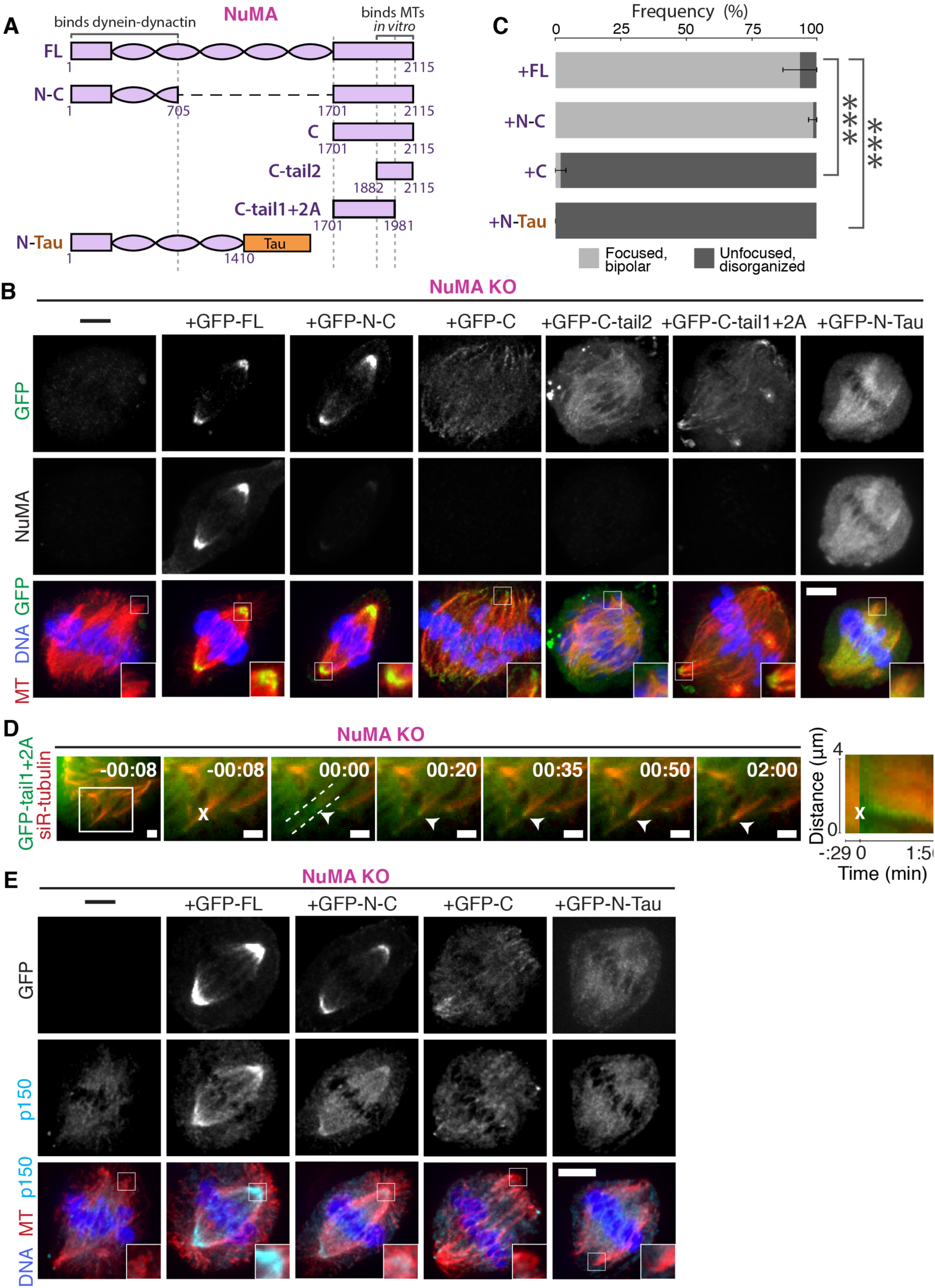
NuMA function requires both minus-end-recognition and dynactin-recruitment modules. See also Table 1, Figure 6-figure supplement 1, and Video 7. (A) Schematic maps of NuMA truncations and chimeras used. ‘FL’ indicates full-length NuMA. ‘N-C,’ ‘C,’‘C-tail1,’ and ‘C-tail2’ are NuMA truncations as indicated. ‘N-Tau’ includes the microtubule binding domain of Tau (orange). Dynein-dynactin bind in the first 705 amino acids of NuMA (Kotak et al., 2012). ‘C-tail2’ was previously implicated in microtubule (MT) binding (Du et al., 2002; Gallini et al., 2016; Haren & Merdes, 2002). (B) Representative immunofluorescence images showing localization of GFP-tagged NuMA truncations and chimeras expressed in RPE1 cells in which endogenous NuMA has been knocked out. Canonical microtubule binding domains in tail2 localize all along spindle microtubules, while the addition of tail1 to tail2A confers minus-end recognition. Note that NuMA antibody does not recognize NuMA’s C-terminus. Scale bar, 5 μm. (C) Graph of the mean percentage of cells ± SEM for each condition from data in (B) that display bipolar spindles with focused poles (light gray) compared to disorganized spindle architecture characteristic of NuMA knockout (detached centrosomes, loss of two focused poles; dark gray). *n* = 19 (‘FL’); *n* = 32 (‘N-C’); *n* = 20 (‘C’); *n* = 16 (‘N-Tau’) from 3-5 independent experiments. \*\*\**P* ≤ 0.0002. (D) Representative time-lapse live images and kymograph of RPE1 cell expressing GFP-Ctail1+2A in a NuMA knockout background. After ablation (‘X’), C-tail1+2A is recruited to k-fiber minus-ends (arrowhead). The k-fiber stub slowly polymerizes, but its minus-end is not transported by dynein and remains detached from the spindle. Time is in min:sec, with the frame captured immediately following ablation set to 00:00 s. Scale bars, 2 μm. Kymograph (right) taken along dashed line path. (E) Representative immunofluorescence images showing localization of dynactin (p150) after NuMA knockout and rescue with GFP-tagged NuMA truncations and chimeras in RPE1 cells. Constructs containing NuMA’s N-terminus recruit p150. Scale bar, 5 μm.

We sought to more closely define which sections within NuMA’s C-terminus (a.a. 1701- 2115) are involved in microtubule minus-end recognition. The second half of the C-terminus (‘Ctail2’, a.a. 1882-2115), which contains residues previously implicated in NuMA-microtubule interactions (Du et al., 2002; Gallini et al., 2016; Haren & Merdes, 2002), localized all along spindle microtubules with no minus-end preference (Figure 6B). Similarly, C-tail2A (a.a. 1882-1981) bound along the lattice, while C-tail1 (a.a. 1701-1881) did not bind microtubules (Figure 6-figure supplement 1). However, in combination (‘C-tail1+2A’, a.a. 1701-1981) they localized at minus-ends (Figure 6B). Indeed, the C-tail1+2A region was sufficient for recruitment to new minus-ends created by ablation, even in the absence of endogenous NuMA (Figure 6D). Thus, minus-end recognition requires amino acids 1701-1881 of the NuMA tail in addition to the lattice-binding C-tail2A. Interestingly, related NuMA residues were recently shown to bind at both plus- and minus-ends and play a role in spindle orientation (Seldin et al., 2016). In sum, our data indicate that NuMA’s localization to spindle microtubule minus-ends is independent of dynein, independent of known direct minus-end binding proteins, and mediated by its C-terminal residues 1701-1981 (‘C-tail1+2A’).

Importantly, NuMA’s C-terminus (‘C’) localized to minus-ends but could not rescue proper pole focusing or spindle architecture, unlike full-length protein or N-C (Figure 6C). This suggests that NuMA’s function in spindle organization requires both minus-end binding (via its C-terminus) and the ability to recruit dynactin to minus-ends (via its N-terminus) (Kotak et al., 2012). Consistent with this hypothesis, NuMA’s C-terminus (‘C’) was unable to recruit dynactin to minus-ends, while its N-terminus and C-terminus fused (‘N-C’) did (Figure 6E; Table 1).

**Table 1.**
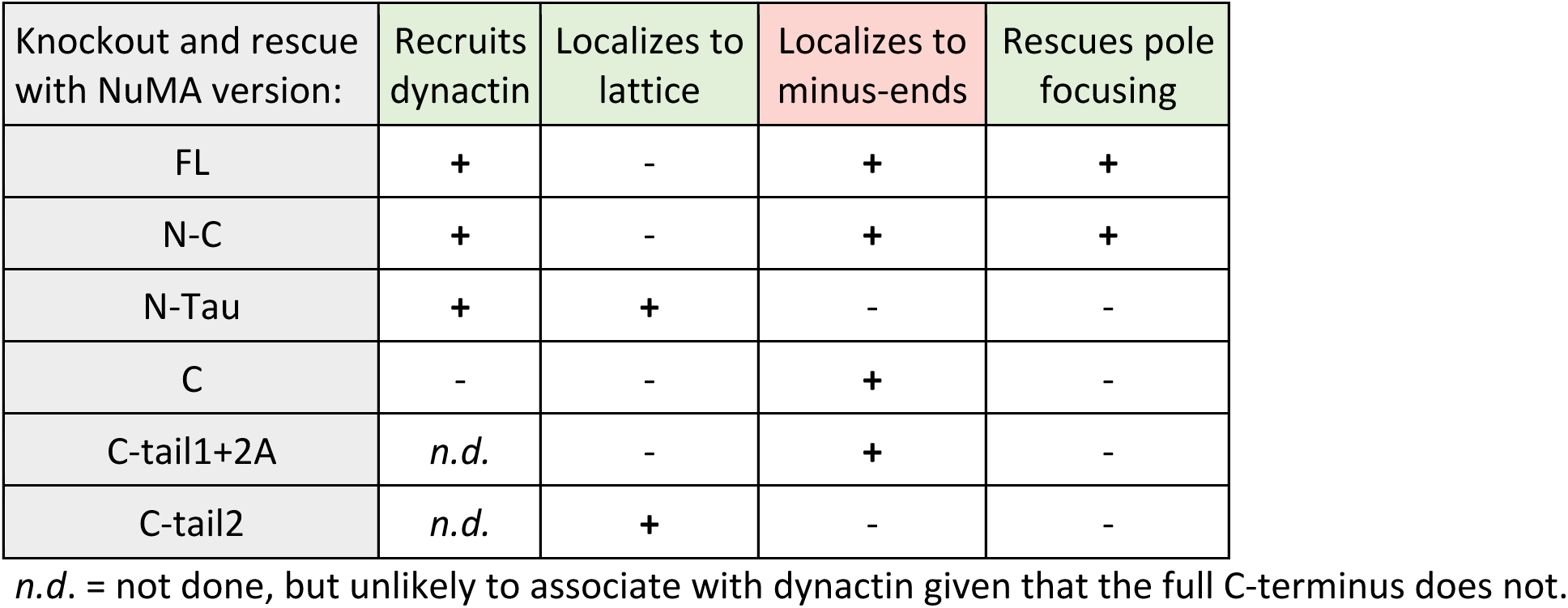
NuMA’s pole-focusing ability correlates with dynactin- and minus-end-binding. See also Figure 6 and Figure 6-figure supplement 1.

| Knockout and rescue with NuMA version: | Recruits dynactin | Localizes to lattice | Localizes to minus-ends | Rescues pole focusing |
| --- | --- | --- | --- | --- |
| FL | + | - | + | + |
| N-C | + | - | + | + |
| N-Tau | + | + | - | - |
| C | - | - | + | - |
| C-tail1+2A | *n.d.* | - | + | - |
| C-tail2 | *n.d.* | + | - | - |

*n.d.* = not done, but unlikely to associate with dynactin given that the full C-terminus does not.

To test whether recruiting dynein-dynactin to the sides of microtubules was sufficient for proper microtubule clustering into poles, we fused NuMA’s N-terminus to the Tau microtubule-binding domain (‘N-Tau’). N-Tau localized along the length of spindle microtubules and recruited dynactin there (Figure 6E). Despite combining dynein-dynactin binding and microtubule lattice binding, N-Tau was not enriched at minus-ends and was unable to rescue spindle architecture (Figure 6C; Table 1).

Like a traditional cargo adaptor, NuMA may target force to spindle minus-ends using a cargo (minus-end) binding module (‘C’) and a dynactin-recruitment module (‘N’). Furthermore, the inability of N-Tau to rescue spindle architecture in the absence of endogenous NuMA suggests that specifically targeting dynein-dynactin to minus-ends, not just all along spindle microtubules as N-Tau does, is critical for organizing a focused, bipolar spindle.

## DISCUSSION

### NuMA targets dynactin to minus-ends, spatially regulating dynein activity at mitosis

Our data indicate that NuMA localizes to spindle microtubule minus-ends independently of dynein and minus-end binding proteins γ-TuRC, CAMSAP1, and KANSL1/3. NuMA then targets dynactin to minus-ends, localizing dynein motor activity there. In addition, targeting dynein to the end of its track could permit amplification by motor pile-up, as NuMA at minus-ends captures processive dynein complexes. Altogether, our findings are consistent with a model in which NuMA confers minus-end targeting of the dynein-dynactin complex upon nuclear envelope breakdown (NEB), when NuMA is released from the nucleus. Indeed, active microtubule clustering by dynein is first observed coincident with NEB (Rusan et al., 2002). Thus, NuMA may provide both spatial (minus-end-specific) and temporal (mitosis-specific) regulation of dynein-powered force (Figure 7A).

**Figure 7.**
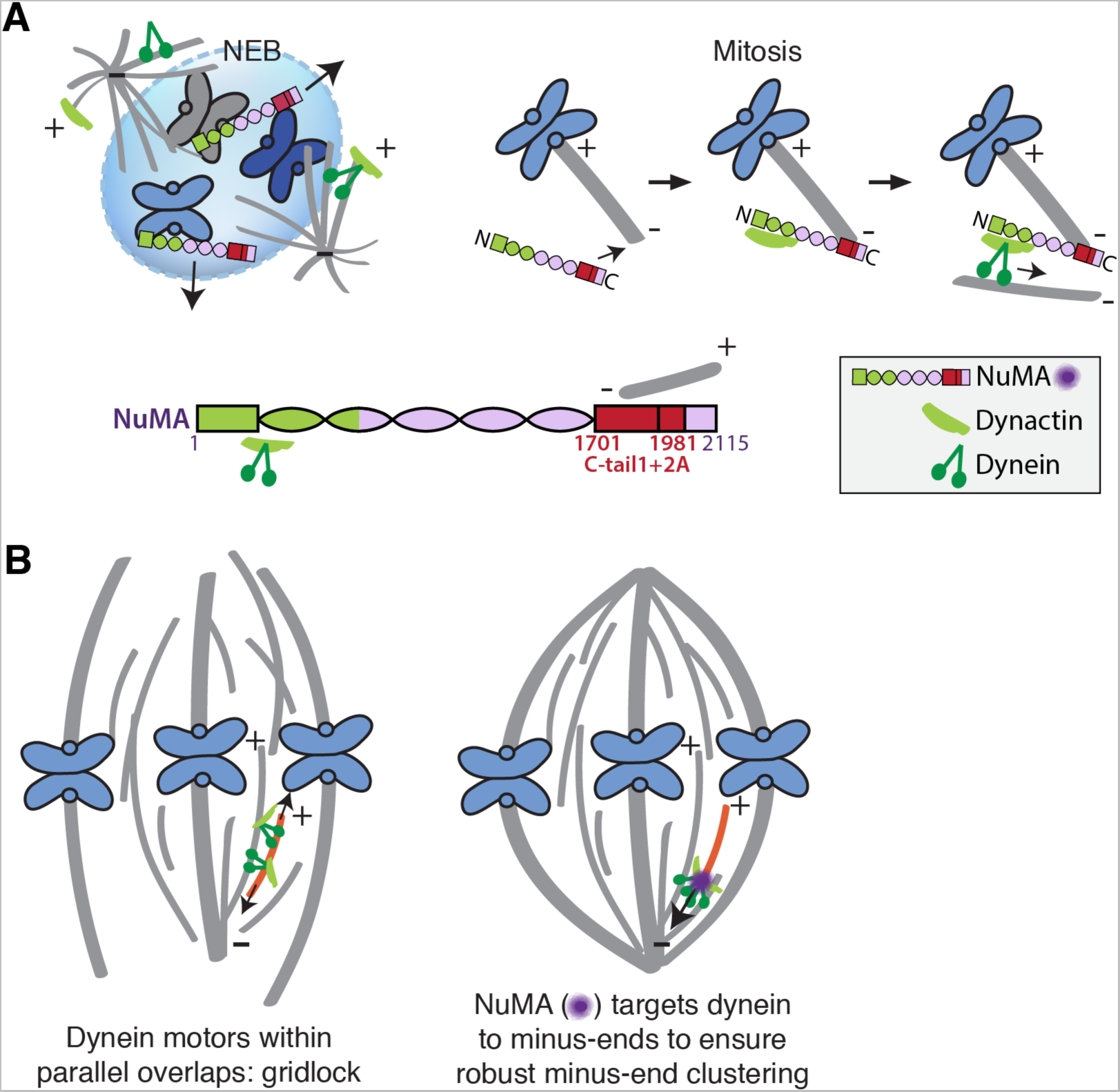
Model: NuMA spatially targets dynein activity to minus-ends at mitosis to ensure minus-end clustering into poles. (A) Our findings suggest that NuMA confers minus-end targeting to the dynein-dynactin complex upon NEB, when NuMA is released from the nucleus. NuMA binding at minus-ends requires a region of its C-terminus (amino acids 1701-1981, in red) that includes tail1 in addition to lattice-binding domain tail2A. NuMA targets dynactin to minus-ends, localizing dynein motor activity. (B) Without minus-end targeting, dynein molecules acting on the red microtubule oppose each other, resulting in gridlock (left). NuMA-mediated targeting of force to minus-ends allows for productive clustering of the red microtubule into the pole (right). Altogether, our data support a model for mammalian spindle organization in which targeting poleward force to microtubule minus-ends specifically – by NuMA-mediated dynactin recruitment – provides robust clustering of microtubules into a focused, bipolar spindle.

The data also indicate that NuMA’s minus-end localization requires residues within its tail1 domain. Previously identified microtubule-binding domains in its C-terminus (tail2A and tail2B) (Du et al., 2002; Gallini et al., 2016; Haren & Merdes, 2002) are not sufficient for minus-end recognition, while tail1+2A is. We propose two hypotheses for tail1+2A-mediated minus-end localization. First, a yet-unidentified minus-end binding protein or the minus-end-directed kinesin HSET (Gaglio et al., 1996; Mountain et al., 1999) could localize NuMA via an interaction that requires NuMA’s tail1 domain. During the preparation of this manuscript, ASPM was identified as a novel mammalian spindle minus-end binder, but ASPM deletion does not affect NuMA localization (Jiang et al., 2017). Second, NuMA tail1+2A may confer minus-end affinity directly *in vivo*, perhaps involving post-translation modifications (Compton & Luo, 1995; Gallini et al., 2016; Magescas et al., 2017; Yan et al., 2015) not previously recapitulated *in vitro*. To robustly bind minus-ends, NuMA could use minus-end-specifying (tail1) and lattice-binding (tail2A) domains in tandem, analogous to the CKK and MBD domains of CAMSAP family proteins (Hendershott & Vale, 2014; Jiang et al., 2014).

### Function of force at spindle microtubule minus-ends

Models of spindle assembly *in silico* predict that enriching a minus-end-directed motor at microtubule ends can break gridlock between parallel microtubules and allow robust minus-end clustering into poles (Burbank et al., 2007; Goshima et al., 2005; Hyman & Karsenti, 1996; Surrey et al., 2001). However, a lack of mechanistic information and tools made it difficult to test such hypotheses *in vivo*. The present study reveals a mechanism for direct recruitment of the motor to minus-ends, via NuMA, and is consistent with the prediction that targeting dynein-dynactin to the sides of microtubules is not sufficient for robust spindle organization. Both the minus-end-binding module (‘C’) and dynein-dynactin-binding module (‘N’) of NuMA are required for bipolar focusing, while the distance between them is not critical for function. Fusing the dynein-dynactin-binding module to the Tau microtubule binding domain is not sufficient, suggesting a requirement for minus-end-specific forces (Figure 6; Table 1). However, we cannot formally exclude that features of the NuMA C-terminus (missing from ‘N-Tau’) other than minus-end binding enable rescue of pole focusing. Altogether, the work is consistent with a model for mammalian spindle organization in which targeting poleward force to microtubule minus-ends (by NuMA-mediated dynactin recruitment) is critical for organizing microtubules into a focused, bipolar metaphase spindle (Figure 7B). More broadly, the emergence of pole architecture at mitosis illustrates how spatial regulation of molecular-scale activities (like NuMA localizing dynein activity at individual minus-ends) can give rise to complex and diverse cellular-scale structures.

### NuMA as mitotic dynein adaptor or activator

The data suggest that NuMA in the mammalian spindle body may function as a traditional cargo adaptor, with a cargo (minus-end)-binding module and a motor-binding module. In the spindle body context, microtubule minus-ends are the cargo, just as the cortex can be framed as cargo for cortical dynein – analogous to more canonical interphase cargo like membranous vesicles or organelles. This framework raises the question of whether NuMA additionally serves as a dynein activator, inducing highly processive motility like interphase dynein adaptors can (McKenney et al., 2014; Schlager et al., 2014).

Dynein’s localization in mammalian mitosis is more ubiquitous than that of dynactin (Figure 4-figure supplement 1; Figure 2-figure supplement 1C); high cytoplasmic levels of dynein prevented us from detecting clear dynein enrichment at ablation-created minus-ends, for example (data not shown), and dynein intermediate chain localization is not as altered by NuMA knockout as dynactin localization (Figure 4-figure supplement 1). These observations could be explained if NuMA and dynactin at minus-ends selectively activate dynein there – through a conformational shift that renders the motor more processive, for example – rather than simply selectively localizing it there. In other words, dynein’s localization within the spindle body may be less tightly regulated than its activity. The loss of dynein activity at minus-ends observed after NuMA knockout (Figure 3C) may stem from a lack of dynein activation (without NuMA and dynactin present at minus-ends) rather than a lack of dynein enrichment.

Unlike known dynein activators, NuMA is thought to not only homodimerize but also oligomerize into higher order assemblies (Harborth et al., 1999; Saredi, Howard, & Compton, 1997). NuMA’s oligomerization ability raises the possibility that it could both activate and assemble teams of motors, similar to adaptors on large cellular cargoes like mitochondria or phagosomes (Fu & Holzbaur, 2014; Rai et al., 2013). The increased force and processivity provided by teams of NuMA-dynactin-dynein complexes on mitotic minus-ends could enable transport and clustering of minus-ends despite high loads and friction created by dense microtubule crosslinking and – in the case of k-fiber minus-ends – coupling to chromosomes.

## MATERIALS AND METHODS

### Cell culture and transfection

PtK2 cells were cultured in MEM (11095; Thermo Fisher, Waltham, MA) supplemented with sodium pyruvate (11360; Thermo Fisher), nonessential amino acids (11140; Thermo Fisher), penicillin/streptomycin, and 10% heat-inactivated fetal bovine serum (FBS) (10438; Thermo Fisher). RPE1 and HeLa cells were cultured in DMEM/F12 with GlutaMAX (10565018; Thermo Fisher) supplemented with penicillin/streptomycin and 10% FBS. For Tet-on inducible CRISPR-Cas9 cell lines, tetracycline-screened FBS (SH30070.03T; Hyclone Labs, Logan, UT) was used. All cells were maintained at 37°C and 5% CO_2_. Cells were transfected with DNA using ViaFect (E4981; Promega, Madison, WI) 48 h (RPE1/HeLa) or 72 h (PtK2) before imaging.

### Inducible CRISPR-Cas9 knockout cells

sgRNAs were designed against 5’ exons of NuMA, CAMSAP1, and KANSL1 using http://crispr.mit.edu. sgRNAs are listed in Table S1. The plasmid used to express sgRNAs under control of the hU6 promoter (pLenti-sgRNA) was a gift from T. Wang, D. Sabatini, and E. Lander (Whitehead/Broad/MIT). An RPE1 cell line containing doxycycline-inducible human codon-optimized spCas9 was a gift from I. Cheeseman (Whitehead/MIT) and was generated as described in (McKinley et al., 2015) using a derivative of the transposon described in (G. Wang et al., 2014). We infected this inducible-spCas9 RPE1 cell line with each pLenti-sgRNA as described in (T. Wang et al., 2015) using virus expressed in HEK293T cells and 10 μg/mL polybrene and selected with 6 μg/mL puromycin. For each targeted gene, we tested 3 independent sgRNA sequences, each of which generated indistinguishable spindle phenotypes (data not shown), and picked one line for subsequent studies. 4 days before each experiment, spCas9 expression was induced with 1 μM doxycycline hyclate.

### Live imaging and laser ablation

For live imaging, cells were plated on glass-bottom 35mm dishes coated with poly-D-lysine (MatTek Corporation, Ashland, MA) and imaged in a stage-top humidified incubation chamber (Tokai Hit, Fujinomiya-shi, Japan) maintained at 30°C and 5% CO2. To visualize tubulin,100 nM siR-Tubulin dye (Cytoskeleton, Inc., Denver, CO) was added 2 h prior to imaging, along with 10 μM verapamil (Cytoskeleton, Inc.). Under these conditions, there was no detected defect in spindle appearance or microtubule dynamics. As described elsewhere (Elting et al., 2014), cells were imaged using a spinning disk confocal inverted microscope (Eclipse Ti-E; Nikon Instruments, Melville, NY) with a 100X 1.45 Ph3 oil objective through a 1.5X lens, operated by MetaMorph (7.7.8.0; Molecular Devices, Sunnyvale, CA). Laser ablation (30 3-ns pulses at 20Hz) with 551 nm light was performed using the galvo-controlled MicroPoint Laser System (Andor, Belfast, UK). For laser ablation experiments, images were acquired more slowly prior to ablation and more rapidly after ablation (typically 7 s prior and 3.5 s after ablation).

### Immunofluorescence and antibodies

For immunofluorescence, cells were plated on #1.5 25 mm coverslips coated with 1 mg/mL poly-L-lysine. Cells were fixed with 95% methanol + 5 mM EGTA at −20°C for 3 min, washed with TBS-T (0.1% Triton-X-100 in TBS), and blocked with 2% BSA in TBS-T for 1 h. Primary and secondary antibodies were diluted in TBS-T + 2% BSA and incubated with cells overnight at 4°C (primary) or for 20 min at room temperature (secondary). DNA was labeled with Hoescht 33342 (Sigma, St. Louis, MO) before cells were mounted in ProLongGold Antifade (P36934; Thermo Fisher). Cells were imaged using the spinning disk confocal microscope described above. Antibodies: mouse anti-α-tubulin DM1α (T6199; Sigma), rabbit anti-α-tubulin (ab18251; Abcam, Cambridge, UK), rabbit anti-NuMA (NB500-174; Novus Biologicals, Littleton, CO), mouse anti-p150-Glued (610473; BD Biosciences, San Jose, CA), mouse anti-α-tubulin DM1α conjugated to Alexa488 (8058S; Cell Signaling, Danvers, MA), mouse anti-dynein intermediate chain (MAB1618MI; Millipore, Billerica, MA), rabbit anti-EB1 (sc-15347; Santa Cruz Biotechnology, Dallas, TX), rabbit anti-KANSL1 (PAB20355; Abnova, Taipei City, Taiwan), rabbit anti-CAMSAP1 (NBP1-26645; Novus Biologicals), rabbit anti-g-tubulin (T3559; Sigma), and camel nanobody against GFP coupled to Atto488 (gba-488; ChromoTek, Hauppauge, NY).

### STORM

PtK2 cells expressing GFP-α-tubulin (gift of A. Khodjakov, Wadsworth Center) were plated on photo-etched, gridded coverslips (G490; ProSciTech, Kirwan, Australia) coated with 1 mg/mL poly-L-lysine (P-1524; Sigma) and imaged at 29-30°C in a homemade heated aluminum coverslip holder using the confocal microscope and ablation system described above. 20-30 s after k-fiber ablation, imaging media was replaced with fixative (as above) chilled to −80°C, and the coverslip holder was placed on ice for 1 min. Cells were incubated with 3% BSA in PBS for 1 hr at RT, and then with primary antibodies overnight at 4°C. Secondary antibodies ((anti-mouse Cy3B; Jackson Immunoresearch, West Grove, PA); anti-rabbit AF647 (Life Tech, Carlsbad,CA)) were incubated for 30 min at RT. Antibody incubations were followed by 4 washes with 0.2% BSA in PBS. Samples were stored in PBS during confocal imaging, and coverslip grid was used to re-find the individual ablated cell. For 3D STORM imaging, samples were mounted on glass slides with a standard STORM imaging buffer consisting of 5% (w/v) glucose, 100 mM cysteamine, 0.8 mg/mL glucose oxidase, and 40 μg/mL catalase in 1M Tris-HCI (pH 7.5) (Huang et al., 2008; Rust et al., 2006). Coverslips were sealed using Cytoseal 60. STORM imaging was performed on a homebuilt setup based on a modified Nikon Eclipse Ti-E inverted fluorescence microscope using a Nikon CFI Plan Apo λ 100x oil immersion objective (NA 1.45). Dye molecules were photoswitched to the dark state and imaged using either 647- or 560-nm lasers (MPB Communications, Montreal, CAN); these lasers were passed through an acousto-optic tunable filter and introduced through an optical fiber into the back focal plane of the microscope and onto the sample at intensities of ~2 kW cm^-2^. A translation stage was used to shift the laser beams towards the edge of the objective so that light reached the sample at incident angles slightly smaller than the critical angle of the glass-water interface. A 405-nm laser was used concurrently with either the 647- or 560-nm lasers to reactivate fluorophores into the emitting state. The power of the 405-nm laser (typical range 0–1 W cm^-2^) was adjusted during image acquisition so that at any given instant, only a small, optically resolvable fraction of the fluorophores in the sample were in the emitting state. Emission was recorded with an Andor iXon Ultra 897 EM-CCD camera at a framerate of 220 Hz, for a total of ~120,000 frames per image. For 3D STORM imaging, a cylindrical lens of focal length 1 m was inserted into the imaging path so that images of single molecules were elongated in opposite directions for molecules on the proximal and distal sides of the focal plane (Huang et al., 2008). Two-color imaging was performed via sequential imaging of targets labelled by AF647 and Cy3B. The raw STORM data was analyzed according to previously described methods (Huang et al., 2008; Rust et al., 2006).

### Drug treatment and microtubule re-growth

To inhibit γ-tubulin, 30 μM gatastatin (gift of Takeo Usui and Ichiro Hayakawa, University of Tsukuba and Okayama University, respectively) (Chinen et al., 2015) was added 25 - 60 min before imaging (from 30 mM stock). For microtubule re-growth (nocodazole washout) experiments, cells were treated with 5 μM nocodazole (M1404; Sigma) for 15 min at 37°C. After 3 washes, cells were incubated at room temperature for 6-8 min before fixation and immunofluorescence (as above).

### Plasmids

2xGFP-Arp1A was made by inserting EGFP from pEGFP-N1 (Clontech, Takara Bio USA, Mountain View, CA) by Gibson assembly between GFP and Arp1A of GFP-Arp1A (human Arp1A in a pBABE variant, Addgene 4432; gift from I. Cheeseman, Whitehead Institute) (Kiyomitsu & Cheeseman, 2012). To make Cas9-resistant GFP-NuMA (‘GFP-NuMA_resistant’), full-length human NuMA (NM_006185.3) with silent mutations (5′-GTGTCAGAGAGACTGGACTTT-3’ mutated to 5′-GTTAGTGAACGCTTGGATTTT-3′, preserving amino acids 57-62 of NP_006176.2 (‘VSERLD’)) was synthesized and cloned (Epoch Life Science, Missouri City, TX) into pEGFP-N1 at BglII and EcoRI sites. Subsequent truncations of NuMA (‘N-C’, ‘C’, ‘C-tail1’, ‘C-tail2’, ‘C-tail2A’, ‘C-tail2B’) were synthesized and cloned (Epoch Life Science) into ‘GFP-NuMA_resistant’ at BglII and HindIII sites. To make GFP-N-Tau, NuMA amino acids 1-1410 from ‘GFP-NuMA_resistant’ followed by a flexible linker and MAPTau (NM_01684.1) from pmEmerald-MAPTau-C-10 (gift from M. Davidson, Florida State University) were synthesized and cloned (Epoch Life Science) into ‘GFP-NuMA_resistant’ at HindIII and SalI sites. Other plasmids used: DsRed-p150^217-548^ (CC1; amino acids 217-548 of chicken p150 in pDsRed-N1, Clontech, gift from T. Schroer, Johns Hopkins University) (Quintyne & Schroer, 2002); mCherry-p50 (chicken p50 in mCherry-C1, Clontech, gift from M. Moffert and T. Schroer, Johns Hopkins University) (Shrum, Defrancisco, & Meffert, 2009); GFP-NuMA (human NuMA in pEGFP-N1, Clontech, gift from D. Compton, Dartmouth Medical School) (Kisurina-Evgenieva et al., 2004); GFP-CAMSAP1 (human CAMSAP1 in pEGFP-C1, Clontech, gift from A. Akhmanova, Utrecht University) (Jiang et al., 2014).

### Data Analysis

To determine the percentage of p150 at plus-ends vs. minus-ends (Figure 1B, Figure 4C), we used single microtubules where both ends were clearly visible, determined p150 localization relative to the EB1-labeled plus-end, and calculated the percentage of p150 at each location within each cell. Percentages for multiple cells were averaged for Figure 1B and 4C. Pre-NEB cells were distinguished from post-NEB cells by the exclusion of microtubules from the nucleus, circle-shaped chromosome packing in the nucleus, and, when possible, NuMA localization within the nucleus.

Kymographs of GFP-Arp1A, GFP-NuMA, and GFP-CAMSAP1 puncta and pole position over time (Figure 1C-E, Figure 2C, Figure 6D) were generated in ImageJ (Version 2.0.0/1.51h). To measure GFP intensity at ablation-created minus-ends over-time (Figure 1F, Figure 2D, Figure 5B), we used a home-written Matlab (R2012a Version 7.4) program to integrate GFP intensity within a 1.4 μm-diameter circle centered on the manually-tracked k-fiber minus-end, and to measure local background intensity within a surrounding 2.7 μm-diameter ‘donut’. After background subtraction, the intensity measured at the cut site during the three frames before ablation (k-fiber intensity) was averaged and set to zero. For NuMA and Arp1A, we then fit a sigmoid function (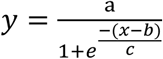, where *y* = intensity and *x* = time) to each trace, normalized to plateau height (*a* = 1), and solved for the time *b* at which *y* = 0.5\**a* to determine time to half-maximum intensity (Figure 1G, Figure 2E, Figure 5C). For CAMSAP1, we normalized each trace to peak height (mean intensity from t = 5 s to t = 20 s) and found the first point at which intensity passed 0.5 to determine time to half-maximum intensity. Finally, to generate mean intensity traces, data from all traces were collected into 5 s wide bins in time and their intensities were averaged. Stub length (distance between k-fiber plus- and minus-ends, Figure 1H) was measured in ImageJ at the first frame following ablation.

Minus-end position data (Figure 3C) were generated by manual tracking of ablation-created k-fiber minus-ends (marked by GFP-CAMSAP1) and spindle poles in time-lapse videos, using a second home-written Matlab program. Nearest neighbor distances between NuMA puncta in STORM imaging (Figure 5H) were measured as the center-to-center distance from each NuMA puncta to its nearest neighboring puncta. NuMA truncation rescue capability (Figure 6C) reports the percentage of bipolar spindles with two focused poles compared to disorganized spindle architecture characteristic of NuMA knockout (detached centrosomes, loss of k-fiber focusing into two poles). Percentage was calculated for each experiment (*n* = 3-5 experiments) and then averaged.

### Statistics

All data are expressed as average ± standard error of the mean (SEM). Calculations of correlation coefficients (Pearson’s *r*) and *p*-values were performed in Matlab. All other *p*-values were calculated using two-tailed unpaired *t*-tests with GraphPad Software. Quoted *n*’s are described in more detail in Figure Legends, but in general refer to individual measurements (biological replicates, e.g., individual spindle lengths, intensity measurements over time after an individual k-fiber ablation, etc.).

### Image presentation

Time-lapse images (Figure 1C-E, 2C, 3A, 4A, 5D, 6D) show a single spinning disk confocal slice, as do immunofluorescence images of individual microtubules (Figure 1A-B, 4C) and post-ablation confocal immunofluorescence images (Figure 5E, Figure 2–figure supplement 1A-B). 3D STORM images (Figure 5F-G) show a single 600 nm slice in Z. Immunofluorescence images of spindles (Figure 2F, 4B, 5A, 6B, 6E and Figure 1–figure supplement 1, Figure 2–figure supplement 1C, Figure 4–figure supplement 1, Figure 6–figure supplement 1B) show max intensity projections (1 – 2 μm in Z) of spinning disk confocal Z-stacks.

### Video preparation

Videos show a single spinning disk confocal Z-slice imaged over time and were formatted for publication using ImageJ and set to play at 35x relative to real time. Videos were corrected to play at a constant frame rate, even when the acquisition rate was not constant.

## ACKNOWLEDGEMENTS

We are grateful to Kara McKinley and Iain Cheeseman for inducible Cas9/CRISPR RPE1 cells, inducible DHC knockout HeLa cells, and helpful advice, and to Anna Akhmanova for GFPCAMSAP1 and sharing unpublished CAMSAP1 data. We thank Takeo Usui and Ichiro Hayakawa for gatastatin, Alexey Khodjakov for PtK2 GFP-α-tubulin cells, and Duane Compton, Iain Cheeseman, Michael Davidson, and Trina Schroer for constructs (GFP-NuMA, GFP-Arp1A, mEmerald-MAPTau, and DsRed-CC1 + mCherry-p50, respectively). Many thanks to Richard McKenney, Ruensern Tan, Tim Mitchison, and the Dumont Lab for discussions and critical reading of the manuscript, and to Jonathan Kuhn and Mary Elting for help with image analysis code. S.D. acknowledges support from NIH DP2GM119177, NIH R00GM09433, the Searle Scholar’s Program, and the Rita Allen Foundation. C.L.H. received support from a NSF Graduate Research Fellowship, a NCI F31 NRSA Predoctoral Fellowship, and a UCSF Moritz Heyman Discovery Fellowship. S.K. and K. X. acknowledge support from NSF under CHE-1554717 and the Pew Biomedical Scholars Award.

## AUTHOR CONTRIBUTIONS

S.J.K. performed STORM imaging (Figure 5F,G) and analyzed STORM data, with guidance from K.X. C.L.H. performed and analyzed all other experiments. C.L.H. and S.D. conceived the project, designed experiments, and wrote the manuscript.

## COMPETING INTERESTS

The authors declare no competing financial or non-financial interests.

## FIGURE SUPPLEMENTS AND LEGENDS

**Figure 1 - figure supplement 1.**
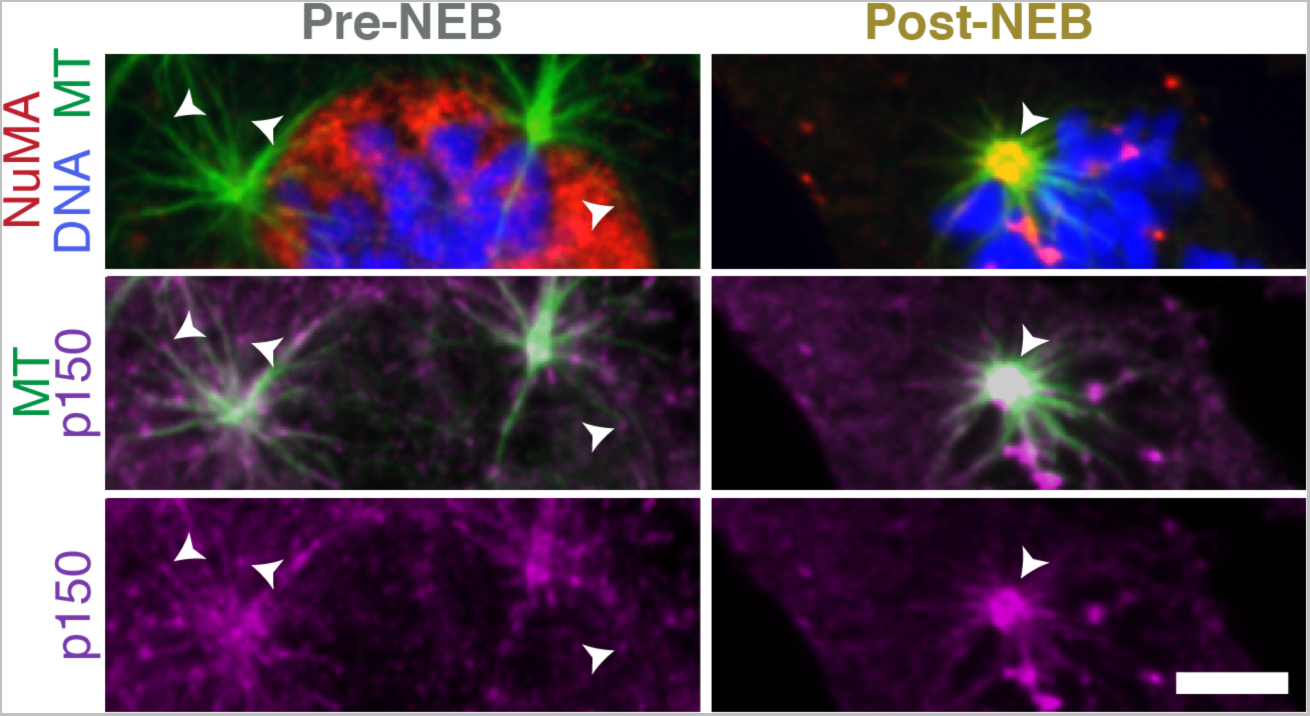
Dynactin localization shifts from plus-ends to minus-ends upon nuclear envelope breakdown. Immunofluorescence showing localization of p150 (dynactin subunit; magenta) and NuMA (red) within asters formed after washout of 5 μM nocodazole in RPE1 cells. Before nuclear envelope breakdown (pre-NEB), NuMA is sequestered in the nucleus and p150 is visible at aster plus-ends (arrowheads), facing outward. After nuclear envelope breakdown (post-NEB), both NuMA and p150 concentrate at aster centers (arrowhead) where minus-ends are. (Similar to Figure 1B, where individual microtubules in the cell periphery with resolvable plus- and minus-ends were analyzed instead of the asters we show here.) Scale bar, 5 μm.

**Figure 2 - figure supplement 1.**
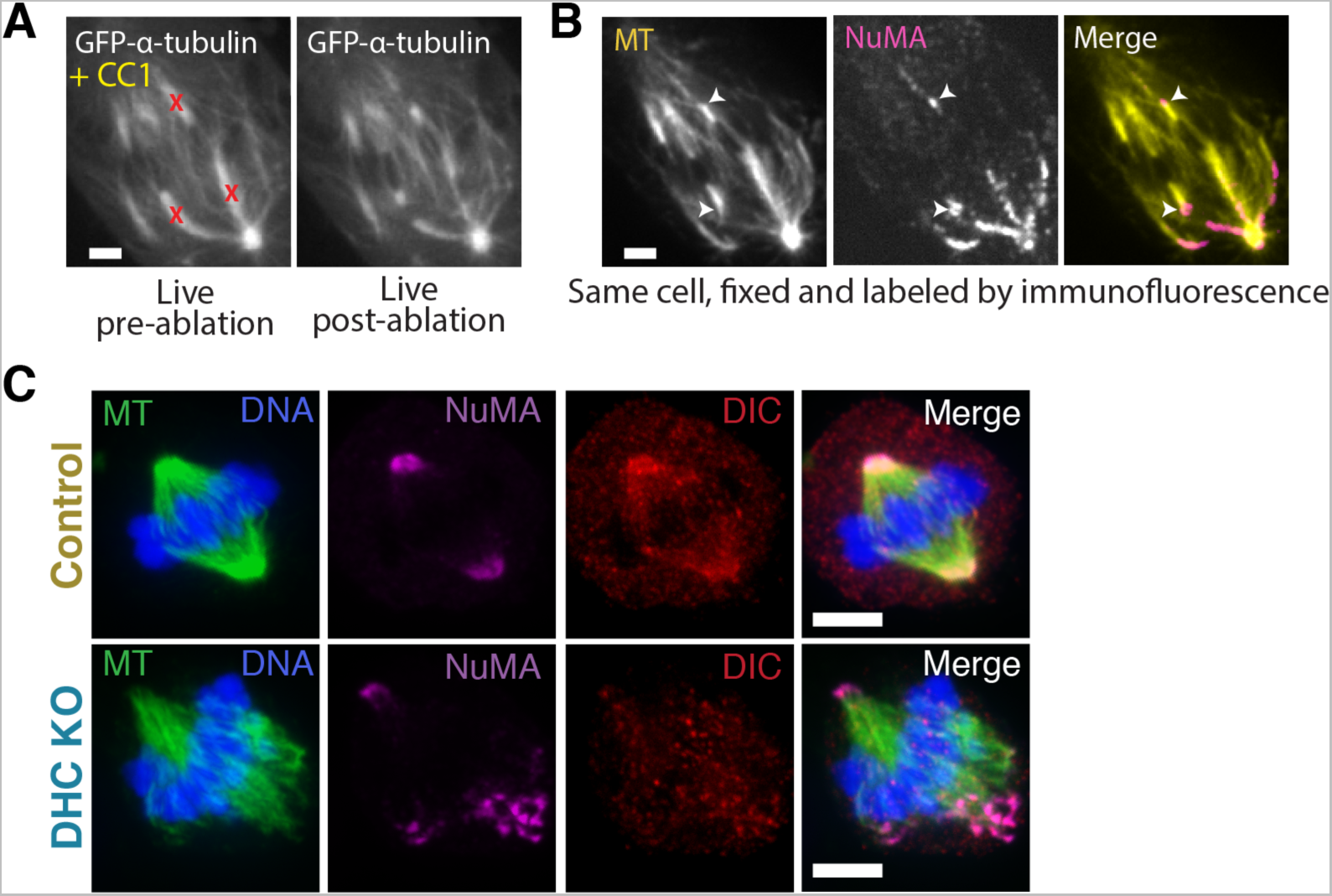
NuMA localizes to minus-ends despite dynein inhibition or knockout. (A) Live images of GFP-α-tubulin immediately before and after k-fiber ablation at targeted sites (red ‘X’s) in a PtK2 cell in which dynein cargo-binding is inhibited by transfection of the dominant negative p150-CC1 fragment (Quintyne & Schroer, 2002). Scale bar, 5 μm. (B) Immunofluorescence image of NuMA (magenta) and α-tubulin (yellow) in cell from (A), fixed after ablation. NuMA (arrowheads) localizes to new minus ends. Scale bar, 5 μm. (C) Immunofluorescence images in inducible-Cas9 dynein heavy chain (DHC)-knockout HeLa cells (McKinley & Cheeseman, 2017) show robust localization of NuMA at minus-ends after DHC knockout. Dynein intermediate chain (DIC) was largely mislocalized after DHC deletion. Scale bar, 5 μm.

**Figure 3 - figure supplement 1.**
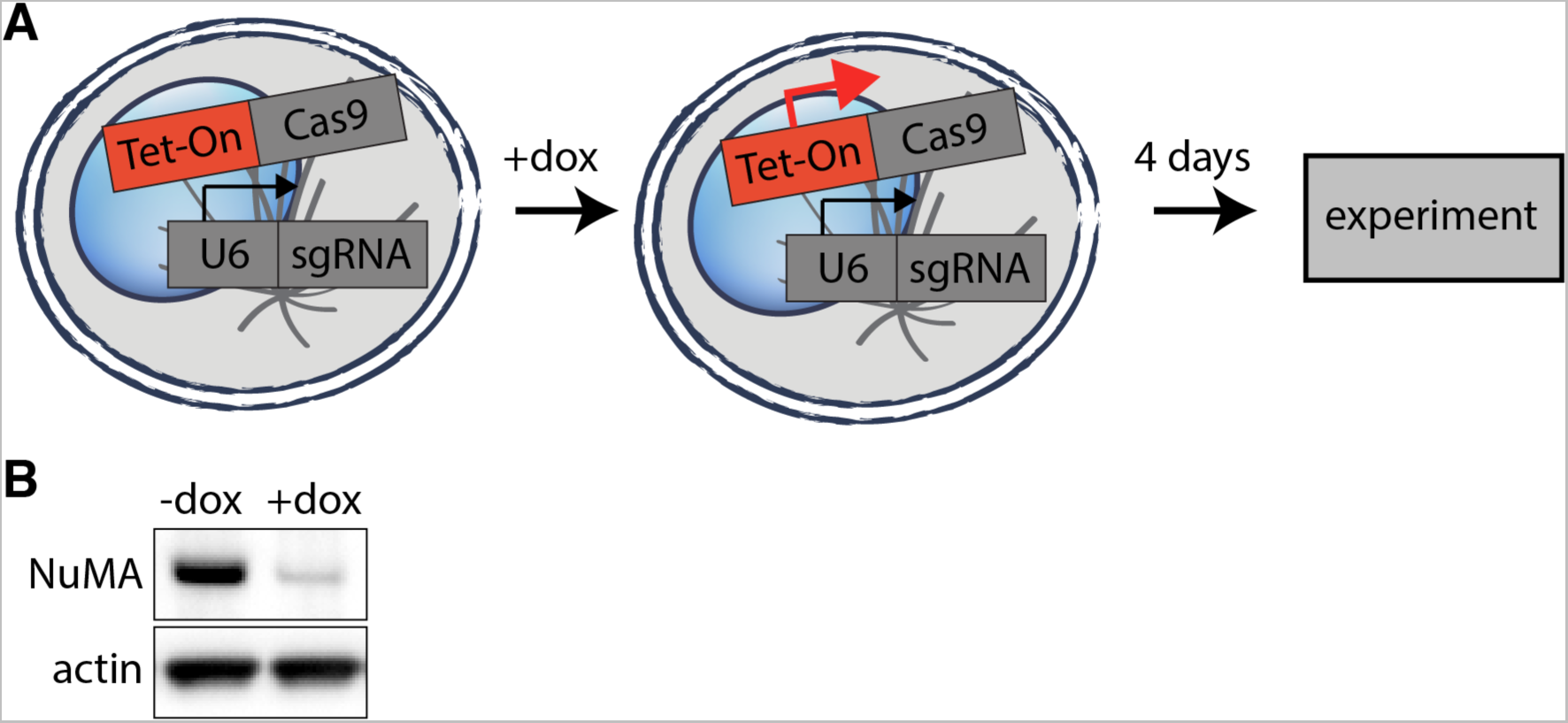
Inducible CRISPR/Cas9 NuMA knockout. (A) Schematic of inducible CRISPR/Cas9 knockout system used (McKinley et al., 2015; T. Wang et al., 2015). Cell lines created have CRISPR/Cas9 and guide RNA stably integrated, but Cas9 expression is only induced upon doxycycline addition 4 days before imaging or analysis, allowing for genetic manipulation of essential mitotic genes. (B) Western blot showing > 90% depletion (normalized to actin) of NuMA protein at the cell population level after knockout. Whenever possible, complete NuMA loss within individual cells analyzed was verified by immunofluorescence.

**Figure 4 - figure supplement 1.**
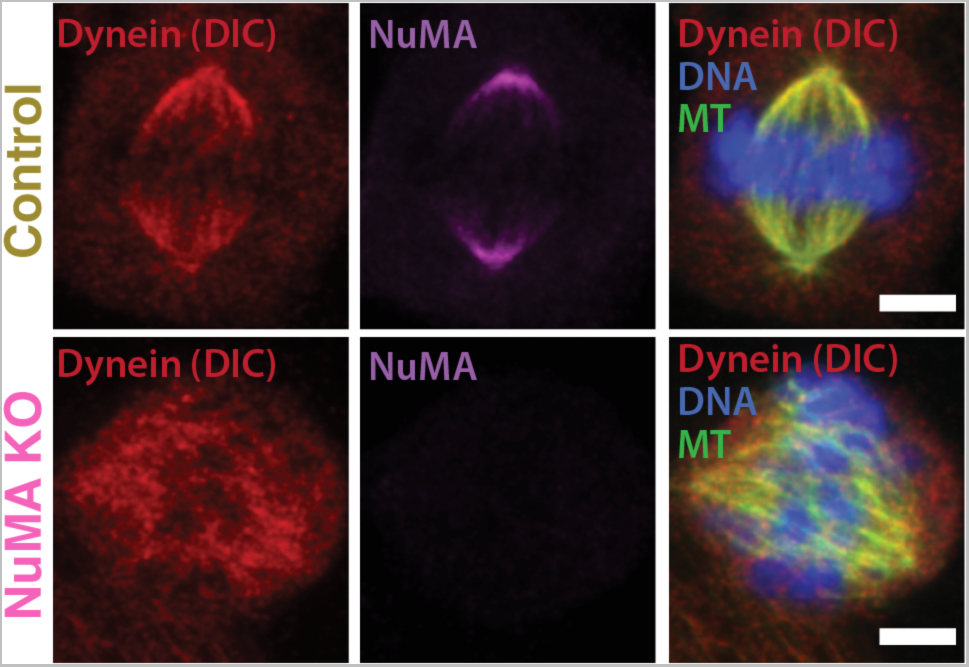
Dynein (DIC) localization after NuMA knockout. Representative immunofluorescence images showing localization of dynein (dynein intermediate chain (DIC); red) and NuMA (magenta) in control RPE1 cells (no doxycycline added) and NuMA knockout cells (after Cas9 induction by doxycycline). Dynein localizes along spindle microtubules in both conditions; its distribution does not appear noticeably altered by NuMA loss (compare to dynactin distribution, Figure 4B). Scale bars, 5 µm.

**Figure 5 - figure supplement 1.**
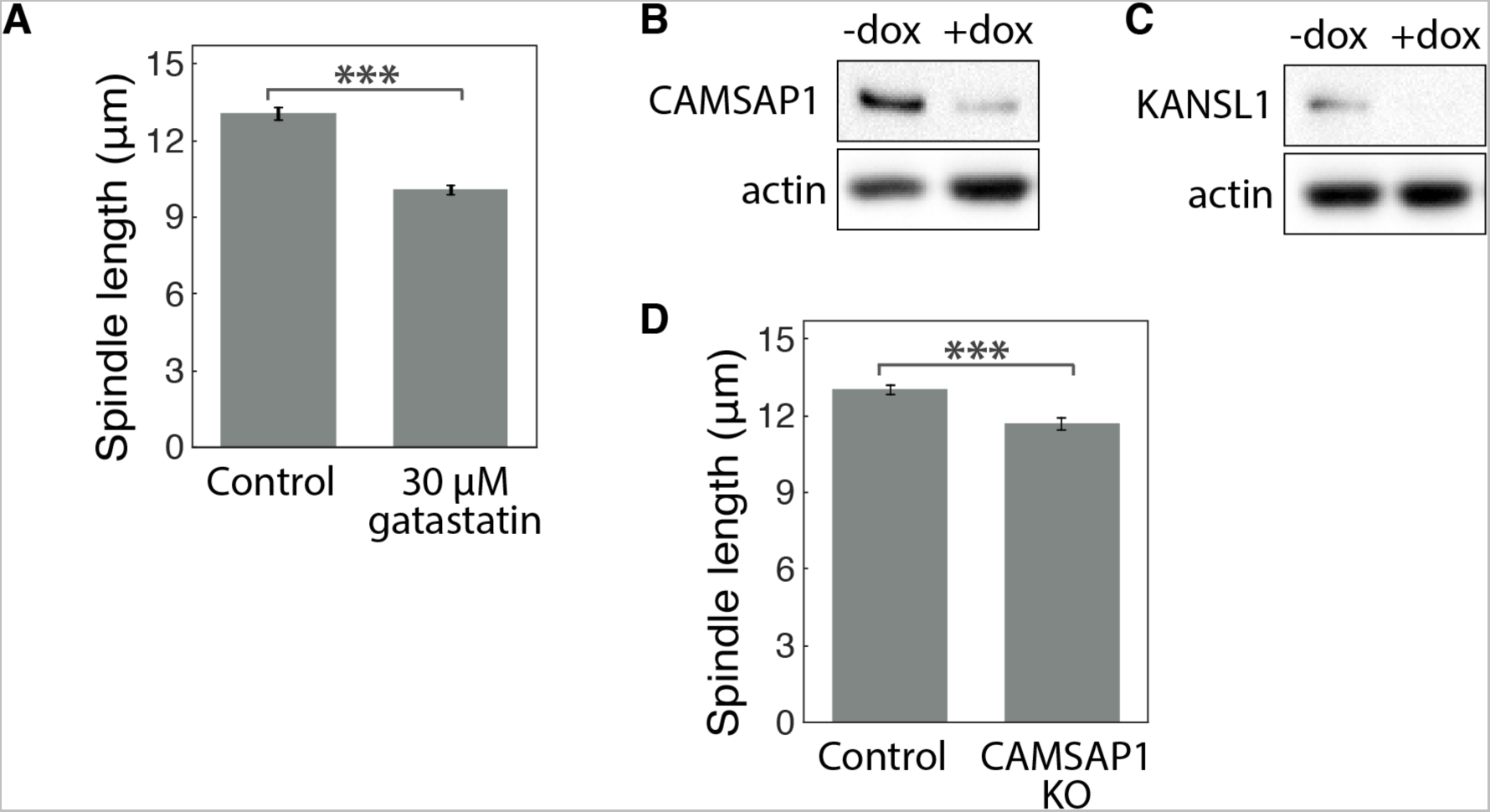
Characterization of gatastatin treatment, CAMSAP1 knockout, and KANSL1 knockout. (A) Mean spindle length ± SEM in control RPE1 cells compared to cells treated with 30 μM gatastatin (γ-tubulin inhibitor) for 25 min. The mean length of gatastatin-treated spindles was reduced by 23%, very similar to previously reported measurements after γ-tubulin inhibition (Chinen et al., 2015). *n* = 24 control cells; *n* = 25 gatastatin-treated cells from 3 separate experiments. \*\*\**P* < 0.0001. (B-C) Western blot showing > 85% depletion (normalized to actin) of (B) CAMSAP1 and (C) KANSL1 protein at the cell population level after knockout (KO). Whenever possible, complete protein loss within individual cells analyzed was verified by immunofluorescence. (D) Mean spindle length ± SEM in control RPE1 cells (no doxycycline added) and CAMSAP1 knockout cells (after Cas9 induction by doxycycline). Complete CAMSAP1 loss was verified by immunofluorescence for all cells analyzed. *n* =11 control cells; *n =*12 CAMSAP1 knockout cells from one experiment. \*\*\**P* < 0.001.

**Figure 6 -figure supplement 1.**
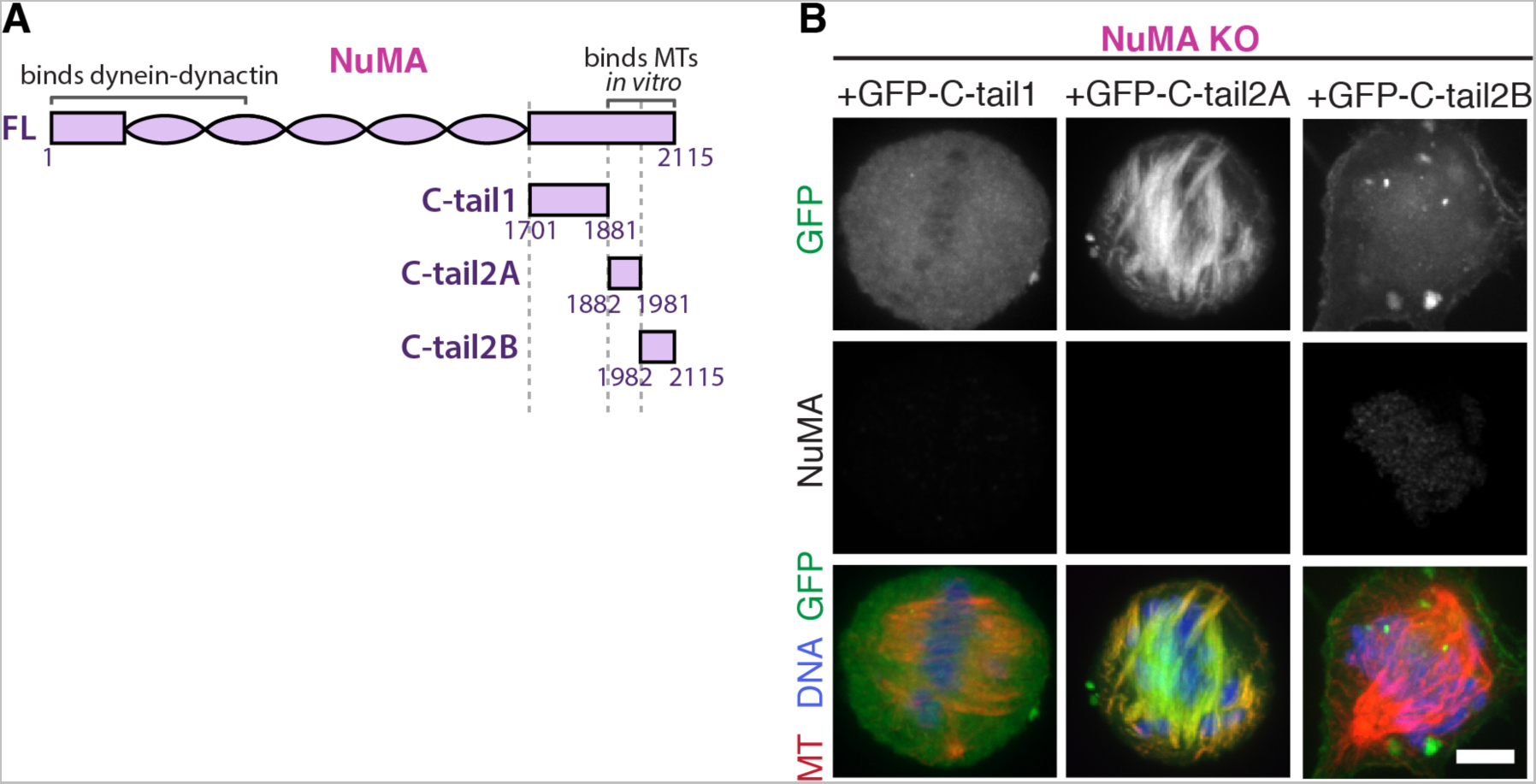
NuMA ‘C-tail1’ and ‘C-tail2A’ alone do not localize to minus-ends. (A) Schematic maps of additional NuMA C-terminal tail truncations. (B) Representative immunofluorescence images showing localization of GFP-tagged NuMA truncations expressed in RPE1 cells in which endogenous NuMA has been knocked out. C-tail1 does not localize to microtubules and C-tail2A localizes all along spindle microtubules, while in combination (‘C-tail1+2A’, Figure 6) they localized at minus-ends. C-tail2B localizes to the cortex but not to microtubules. Scale bar, 5 μm.

**Supplementary Table 1.**
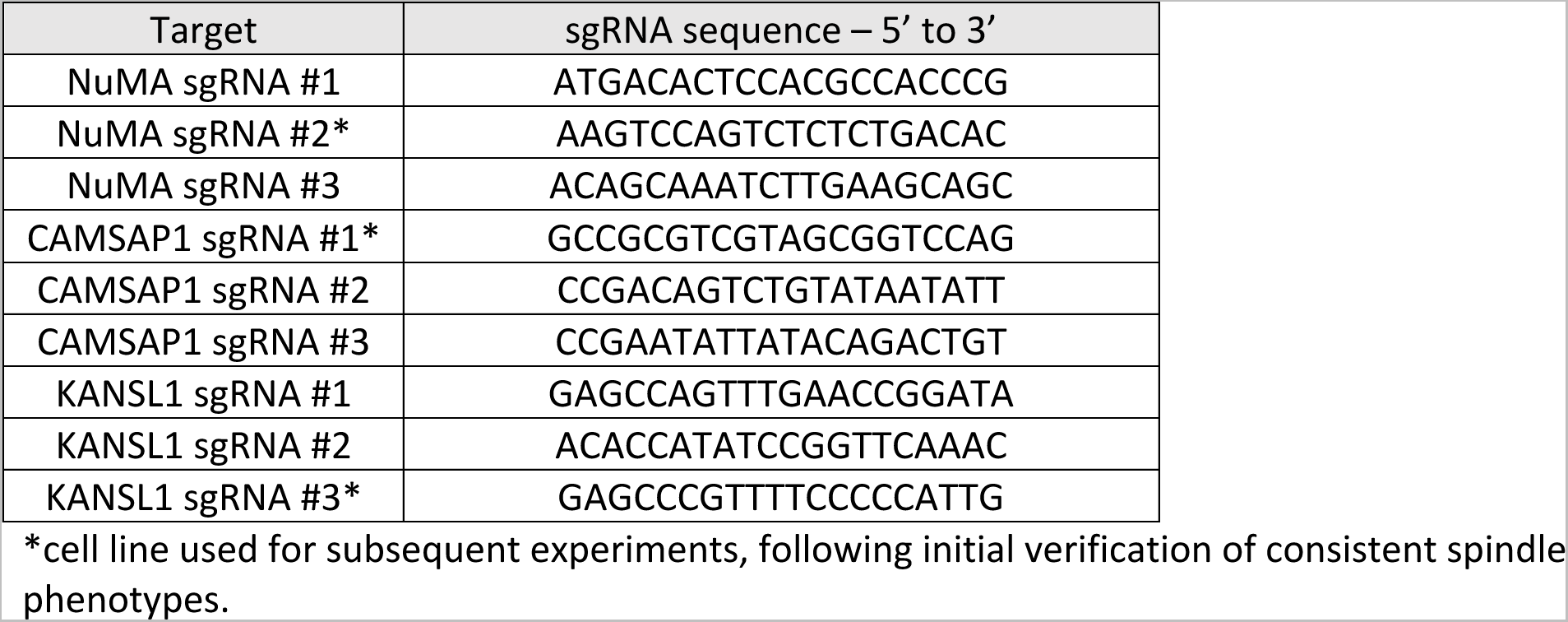
CRISPR-Cas9 sgRNA sequences. See also Materials and Methods.

## VIDEO LEGENDS

**Video 1. Dynactin is robustly recruited to new spindle microtubule minus-ends.** See also Figure 1C.

Live confocal imaging of a PtK2 cell expressing GFP-Arp1A, with spindle microtubules labeled by siR-tubulin. GFP-Arp1A is recruited to k-fiber minus-ends (arrowhead) created by ablation (‘X’) and moves with them as dynein pulls them poleward (Elting et al., 2014). Time is in min:sec, with the frame captured immediately following ablation set to 00:00. Plus-end of ablated k-fiber is marked by ‘*’. Scale bar, 5 μm.

**Video 2. NuMA is robustly recruited to new spindle microtubule minus-ends.** See also Figure 1D.

Live confocal imaging of a PtK2 cell expressing GFP-NuMA, with spindle microtubules labeled by siR-tubulin. GFP-NuMA is quickly recruited to k-fiber minus-ends (arrowhead) created by ablation (‘X’) and moves with them as dynein pulls them poleward (Elting et al., 2014). Time is in min:sec, with the frame captured immediately following ablation set to 00:00. Plus-end of ablated k-fiber is marked by ‘*’. Scale bar, 5 μm.

**Video 3. CAMSAP1 is robustly recruited to new spindle microtubule minus-ends.** See also Figure 1E.

Live confocal imaging of a PtK2 cell expressing GFP-CAMSAP1, with spindle microtubules labeled by siR-tubulin. GFP-CAMSAP1 is immediately recruited to k-fiber minus-ends (arrowhead) created by ablation (‘X’) and moves with them as dynein pulls them poleward (Elting et al., 2014). Time is in min:sec, with the frame captured immediately following ablation set to 00:00. Plus-end of ablated k-fiber is marked by ‘*’. Scale bar, 5 μm.

**Video 4. NuMA quickly localizes to k-fiber minus-ends despite dynein-dynactin inhibition.** See also Figure 2C.

Live confocal imaging of a PtK2 cell expressing GFP-NuMA, in which dynein-dynactin is inhibited by overexpression of p50 (dynamitin) (Shrum et al., 2009). K-fibers are unfocused and splayed due to dynein-dynactin inhibition, but NuMA is still robustly recruited to k-fiber minus-ends (arrowheads) created by ablation (‘X’). Time is in min:sec, with frame captured immediately following ablation set to 00:00. Plus-end of ablated k-fiber is marked by ‘*’. Scale bar, 5 μm.

**Video 5. NuMA is required for transport of minus-ends by dynein.** See also Figure 3A.

Live confocal imaging of a RPE1 cell in which NuMA has been knocked out using an inducible Cas9 system, and GFP-CAMSAP1 is expressed to label minus-ends. After ablation (‘X’), the k-fiber minus-end (arrowhead) is not transported poleward by dynein and remains detached from the disorganized spindle. Time is in min:sec, with the frame captured immediately following ablation set to 00:00 s. Plus-end of ablated k-fiber is marked by ‘*’. Scale bar, 5 μm.

**Video 6. Dynactin is not recruited to minus-ends in the absence of NuMA.** See also Figure 4A. Live confocal imaging of a RPE1 cell in which NuMA has been knocked out using an inducible Cas9 system. In the absence of NuMA, 2xGFP-Arp1A does not localize to k-fiber minus-ends, and its recruitment is not detectable at new minus-ends (arrowheads) created by ablation (‘X’). Note that mislocalized 2xGFP-Arp1A puncta diffuse randomly within the spindle, including within microns of the new minus-end, but do not remain there. Time is in min:sec, with frame captured immediately following ablation set to 00:00. Plus-end of ablated k-fiber is marked by ‘*’. Scale bar, 5 μm.

**Video 7. The** ‘**tail1+2A’ region of NuMA’s C-terminus is sufficient for minus-end localization.** See also Figure 6D.

Live confocal imaging of a RPE1 cell expressing GFP-C-tail1+2A in a NuMA knockout background. After ablation (‘X’), GFP-C-tail1+2A is recruited to k-fiber minus-ends (arrowhead). The k-fiber stub slowly polymerizes, but its minus-end is not transported by dynein and remains detached from the spindle. Note that C-tail1+2A does not rescue spindle architecture; video shows one confocal slice of a three-dimensionally disorganized spindle. Also note that bright GFP signal from a neighboring interphase cell is noticeable on the left side of the video. Time is in min:sec, with the frame captured immediately following ablation set to 00:00 s. Plus-end of ablated k-fiber is marked by ‘*’. Scale bar, 5 µm.

